# Loss of B cell tolerance at the T2/T3a B cell transition is a convergent pathogenic mechanism in common variable immunodeficiency

**DOI:** 10.1101/2025.06.07.658167

**Authors:** Kirsty Hillier, Grace Yuen, Anson Hui, Suparna Kumar, Priyamvada Guha Roy, Joseph T. McColgan, Kimberly Zaldana, Hugues Allard-Chamard, Katie Premo, Naoki Kaneko, Nicole Ingram, Sara Barmettler, Musie Ghebremichael, Jolan E. Walter, Cory A. Perugino, Jishnu Das, Jocelyn R. Farmer, Shiv Pillai

## Abstract

Many patients with common variable immunodeficiency (CVID), including those with CTLA4 deficiency, *NFKB1* variants and activated PI3K-delta syndrome (APDS), develop autoimmunity that is refractory to treatment. Despite this shared clinical phenotype, a unifying mechanism for the breakdown of B cell tolerance across monogenic forms of CVID has not been established. Here, we demonstrate that patients with loss-of-function *NFKB1* variants, like those with *CTLA4* variants and APDS, exhibit dysregulated CD4^+^ T cell expansion, accumulation of transitional B cells, and a relative lack of follicular B cells. In patients with monogenic CVID and clinical autoimmunity, we observed a relative expansion of transitional and activated naïve (aN: CD21^lo^CD11c^hi^) B cells in peripheral blood accompanied by a marked increase in the frequency of VH4-34 expressing autoreactive 9G4^+^ B cells, which expanded between T2 and T3a transitional B cell stages. Single-cell transcriptomic and B cell receptor analysis further revealed a marked expansion of activated T1/2, T3 and extrafollicular activated naïve and double negative (DN: IgD^-^CD27^-^) B cell subsets in APDS patients. Notably, one B cell subset appeared exclusively in the APDS disease state, characterized by high oxidative phosphorylation in transitional B cells, specifically. In APDS patients, we also observed a clonal expansion of specific extrafollicular class-switched DN B cells, which were clonally derived from activated transitional B cells. DN B cells were also identified in APDS lung tissue, consistent with the contribution of activated, extrafollicularly-derived B cells to tissue inflammation. Together, these findings suggest that in many patients with CVID and autoimmune features, premature activation of autoreactive transitional T2 and T3a B cells induces the survival and expansion, rather than the tolerization and elimination, of self-reactive B cells. This process leads to extrafollicular expansion of autoreactive B cells capable of tissue infiltration.

**One Sentence Summary:** Loss of transitional B cell tolerance and extrafollicular expansion of autoreactive B cells drive autoimmunity in monogenic causes of CVID.

## INTRODUCTION

Transitional B cells are the first B cells to emerge from the bone marrow and include a large fraction of self-reactive cells that are culled as they differentiate into mature B cells. Self-reactive B cells that have not been tolerized by receptor editing in the bone marrow or by deletion at transitional B cell stages can be induced by soluble self-antigens in the periphery to enter an anergic state. Peripheral B cell tolerance is maintained by regulatory T cells that restrain T cell help against many self-proteins specifically recognized by anergic B cells. However, anergic B cells may contribute to pathogen responses and induce transient, cross-reactive anti-self-responses [1]. These B cells can also undergo somatic hypermutation and participate in immune responses against foreign antigens, a process sometimes referred to as clonal redemption [2]. In some circumstances, presumably because of the strong antigenic similarity between a self-epitope and a viral antigenic epitope, somatic hypermutation can generate pathogenic autoimmune clones, as described in multiple sclerosis [3].

Inborn errors of immunity (IEI) are a heterogeneous collection of disorders caused by single gene defects in the human immune system that contribute to immune dysregulation. Common variable immunodeficiency (CVID) is the most frequent clinical presentation of IEI with an incidence of 1:25,000 to 1:100,000 [4]. A diagnosis of CVID requires the demonstration of low antibody levels in combination with failed humoral immune response to vaccination.

Immune dysregulation leading to autoimmunity and lymphoproliferation are recognized in upwards of 71% of patients with CVID followed at large tertiary care centers [5–7]. Immune cytopenias are highly prevalent in these patients with a 120-fold higher risk than the general population [5, 6, 8–10]. Furthermore, autoimmune disease portends a higher risk of mortality in CVID than in those with an infectious-only phenotype [5, 11], highlighting the unmet need of identifying pathogenic mechanisms underlying these disease states.

We previously reported a block in B cell development from the transitional to follicular B cell stage in patients with activated PI3K-delta syndrome (APDS), a monogenic form of CVID in which hyperactivation of the mTORC1 pathway results in humoral immunodeficiency, hyperactivation of T cells, and autoimmunity [12]. Patients with APDS are known to have increased activated effector T cells in combination with an expansion of early transitional B cells [13–18], leading us to hypothesize that dysregulated or premature T-B cell collaboration may contribute to a break in self-tolerance at the transitional B cell stage in humans. Our subsequent studies on patients with CTLA4 deficiency strengthened this hypothesis, revealing a similar expansion of transitional B cells in linear correlation with the underlying degree of regulatory T cell dysfunction and effector CD4^+^ T cell expansion [19]. Here we also show that patients with *NFKB1* variants demonstrate excess CD4^+^ T cell activation and T regulatory cell dysfunction paired with a similar accumulation of early T1/2 and T3a type transitional B cells. We further combine detailed flow cytometry along with transcriptomic and B cell receptor profiling to study autoreactive B cells across these three monogenic causes of CVID. Together, our data reveal a marked and convergent expansion of autoreactive transitional and extrafollicular B cells in patients with CVID who present with clinical autoimmunity with further demonstration of tissue-infiltrating activated extrafollicular B cells. These data expand our understanding of convergent mechanisms that drive clinical autoimmunity in CVID.

## RESULTS

### Effector CD4^+^ T cells and early transitional B cell expand in monogenic CVID with autoimmune features

We have previously shown that patients with CTLA4 deficiency and APDS have increased transitional and activated naïve B cells [12, 19, 20]. We now investigated whether this B cell phenotype was also present in another cohort of patients with CVID, namely patients with loss-of-function *NFKB1* variants. We first established that the *NFKB1* variants from the patients included in this study were loss-of-function variants by expressing wild type and variant *NFKB1* gene constructs in the Jurkat human T cell line, and the Sleeping Beauty Transposase ensured integration of the transfected genes. Overexpressed *NFKB1* activity (or the loss of activity) was assessed by transcriptional readouts (see **Methods** and **Supplemental Figure 1** for details).

In patients with loss-of-function *NFKB1* variants, there was a relative accumulation of B cells in the naïve IgD^+^CD27^-^ gate, with very few cells in the marginal zone (IgD^+^CD27^+^) and switched memory (IgD^-^CD27^+^) gates, although some patients had a slight relative increase in double-negative (IgD^-^CD27^-^) B cells. We observed a convergent increase in transitional B cells, including T1/2 (CD10^+^MitoTracker^+^), T3a (CD10^-^MitoTracker^+^CD45RB^-^CD73^-^), and activated naïve B cells (CD21^lo^CD11c^hi^) in patients with CTLA4 deficiency, NFKB1 deficiency, and APDS relative to healthy controls (**Figure 1 and Supplemental Figures 2 and 3**).

**Fig 1.**
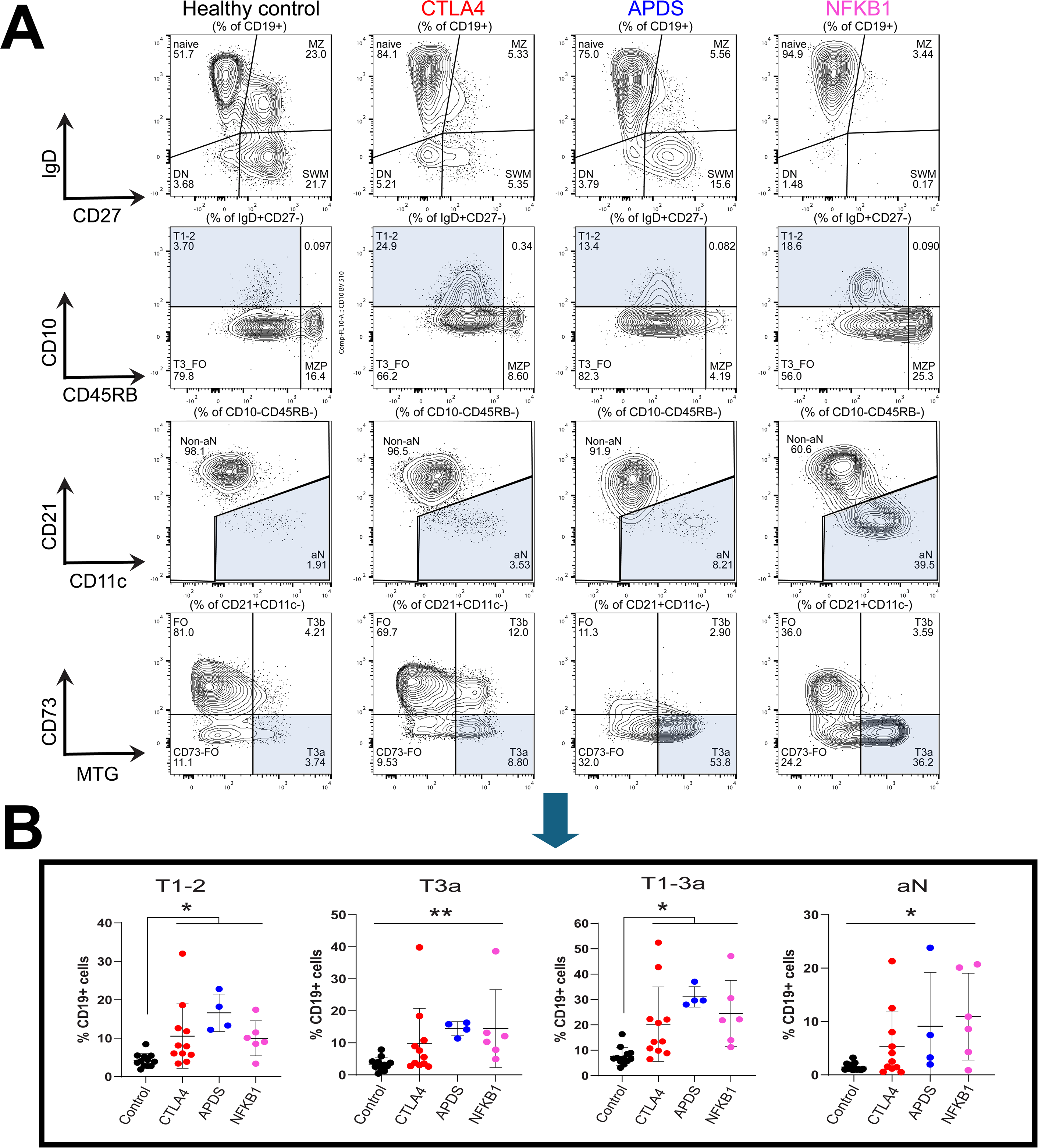
Convergent expansion of early transitional and activated naïve (aN) B cell subsets across monogenic CVID disorders that cause clinical autoimmunity. Flow cytometric analysis of peripheral blood B cell populations in patients with CVID showing (A) flow cytometry plots of representative healthy control and patients with CTLA4 deficiency, APDS, and NFKB1 deficiency, and (B) flow cytometric analysis and quantification of B cells subsets are shown for patients with CTLA4 deficiency (n=11), APDS (n=4), NFKB1 deficiency (n=6), including T1/2 (CD10^+^MitoTracker^+^), and controls (n=12), including transitional T3a (CD10^-^ MitoTracker^+^CD45RB^-^CD73^-^), and activated naïve (aN) (CD21^lo^CD11c^hi^) B cell subsets as percent of CD19^+^ cells. Symbols represent unique individuals; bars represent geometric means with 95% CI; *P < 0.05; **P < 0.01 by Kruskal-Wallis test for non-parametric data

In CTLA4 deficiency and APDS, there is evidence for CD4^+^ T cell dysregulation, demonstrated by the high functional ratio between CD4^+^ effector and regulatory T cells [12, 20]. We hypothesized that NFKB1 may function in a cell-intrinsic manner in T cells to influence regulatory T cells, or alternatively, may directly influence the development or expansion of effector CD4^+^ T cells. We used a CRISPR/Cas9 approach on purified naïve primary human CD4^+^ T cells to functionally inactivate *NFKB1*. *NFKB1-*directed guide RNA induced a marked reduction of NFKB1 protein expression in electroporated T cells. Loss of NFKB1 protein had no effect on *in vitro* regulatory T cell differentiation, but NFKB1-deficient T cells exhibited an increase in proliferation in response to anti-CD3 and anti-CD28 (**Supplemental Figure 4**). These *in vitro* data suggested that the functional loss of NFKB1 may contribute to the expansion of CD4^+^ effector/memory T cells *in vivo*.

To examine whether patients with loss-of-function *NFKB1* variants present with a relative expansion of effector/memory CD4^+^ T cells, we used multiparametric flow cytometry on PBMC samples from these patients. We noted that patients with loss-of-function *NFKB1* variants exhibit a loss of naive (CD62L^+^CD45RA^+^) T cells, and an expansion of central memory (TCM; CD62L^-^CD45RA^-^) and effector memory (TEM; CD62L^+^CD45RA^-^) T cells (**Supplemental Figure 5**). Overall, these data suggest that loss of functional *NFKB1 in vivo* contributes to the expansion of CD4^+^ effector cells, without affecting regulatory T cells. Taken together, these data define a relative expansion of early transitional B cells, convergent across three different monogenic forms of CVID. In all three cohorts of patients, those with APDS or with variants in *CTLA4* or *NFKB1*, there is an increased expansion of CD4^+^ effector T cells relative to regulatory T cells.

### Convergent loss of B cell tolerance at the T1/2 to T3a transition in monogenic CVID with autoimmune features

To define the B cell developmental stage at which tolerance is compromised, we quantitated 9G4 positive B cells in CVID patients with and without clinical autoimmunity. The 9G4 monoclonal antibody identifies a hydrophobic idiotope encoded in the germline-encoded framework region of the immunoglobulin heavy chain (IgH) VH4-34 gene, which is known to have intrinsic autoreactive properties, making 9G4 a useful marker for autoreactivity. This marker has been previously used in both systematic lupus erythematosus and CVID [21–24]. The CVID patient subgroups including APDS (n=4), CTLA4 deficiency (n=11), and NFKB1 deficiency (n=5) and are detailed in the Methods and **Supplemental Table 1**. **Supplemental Figure 2** provides the gating strategy used to define IgD^+^CD27^-^ B cell subsets (T1/2, T3a, and T3b transitional B cell stages, Follicular (FO) B cells, activated Naïve (aN) B cells, and Marginal Zone Precursor (MZP) B cells), as well as switched memory (SWM) B cells, Marginal Zone (MZ) B cells and Double Negative (DN) B cells.

We observed an increased frequency of 9G4^+^ B cells in CVID patients with clinical autoimmunity compared to those without clinical autoimmunity, which markedly expanded at the transitional T3a B cell stage (geometric mean 9G4 positivity 5.4 vs 11% in T1/2 vs T3a B cells from CVID patients with autoimmunity, p<0.001) (**Figure 2A**). These data suggested a break in tolerance, occurring only in the context of clinical autoimmunity and specifically as B cells transitioned from T2 to T3a stages. Further comparing CVID patients with and without autoimmunity, an increased 9G4^+^ B cell frequency was maintained in CVID patients with autoimmunity at the T3b B cell stage (9G4 positivity in T3b B cells 11.3 vs 4.7% in CVID patients with vs without autoimmunity, p=0.005) and in downstream B cell subsets that included aN, SWM, MZP, and MZ (**Figure 2A**). Accounting for differences in the relative frequency of B cell subsets between these patient groups (**Supplemental Figure 6**), these findings were recapitulated with 9G4 positivity evaluated as a percentage of CD19^+^ total B cells (**Supplemental Figure 7**). Overall, these data were consistent with ongoing differentiation of 9G4^+^ B cells, expanding through transitional T3a and T3b B cell stages, in CVID patients with autoimmunity.

**Fig 2.**
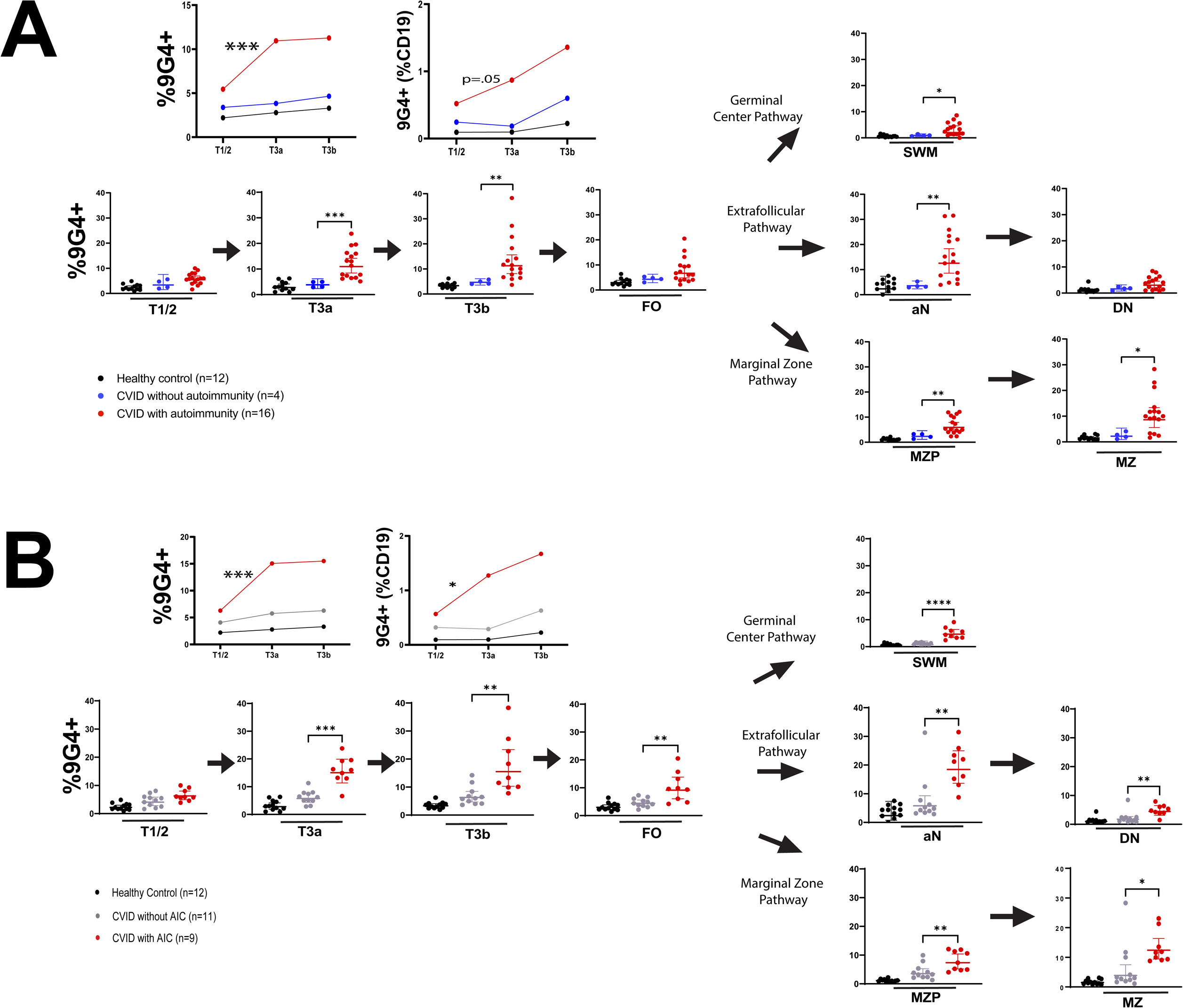
Circulating 9G4^+^ B cell frequency increases between the T2 and T3 B cells stages in patients with common variable immunodeficiency (CVID) and autoimmune clinical manifestations. (A) Flow cytometric analysis of peripheral blood B cell populations in patients with CVID with (n=16) and without autoimmunity (n=4) and healthy controls (n=12), and (B) flow cytometric analysis of peripheral blood B cell populations in patients with CVID with (n=9) and without AIC (n=11) and healthy controls (n=12). Analysis and quantification of B cell subsets are shown, including transitional T1/2 (CD10^+^MitoTracker^+^), transitional T3a (CD10^-^ MitoTracker^+^CD45RB^-^CD73^-^), transitional T3b (CD10^-^MitoTracker^+^CD45RB^-^CD73^+^), follicular (FO) (MitoTracker^-^CD45RB^-^CD73^+^), activated naïve (aN) (CD21^lo^CD11c^hi^), switched memory (SWM) (IgD^-^CD27^+^), marginal zone progenitor (MZP) (CD21^-^CD45RB^+^), marginal zone (MZ) (IgD^+^CD27^+^), and double negative (DN) (IgD^-^CD27^-^). Symbols represent unique individuals; bars represent geometric means with 95% CI; *P < 0.05; **P < 0.01; ***P < 0.001; ****P < 0.0001 by Mann-Whitney U test for non-parametric data.

**Fig 3.**
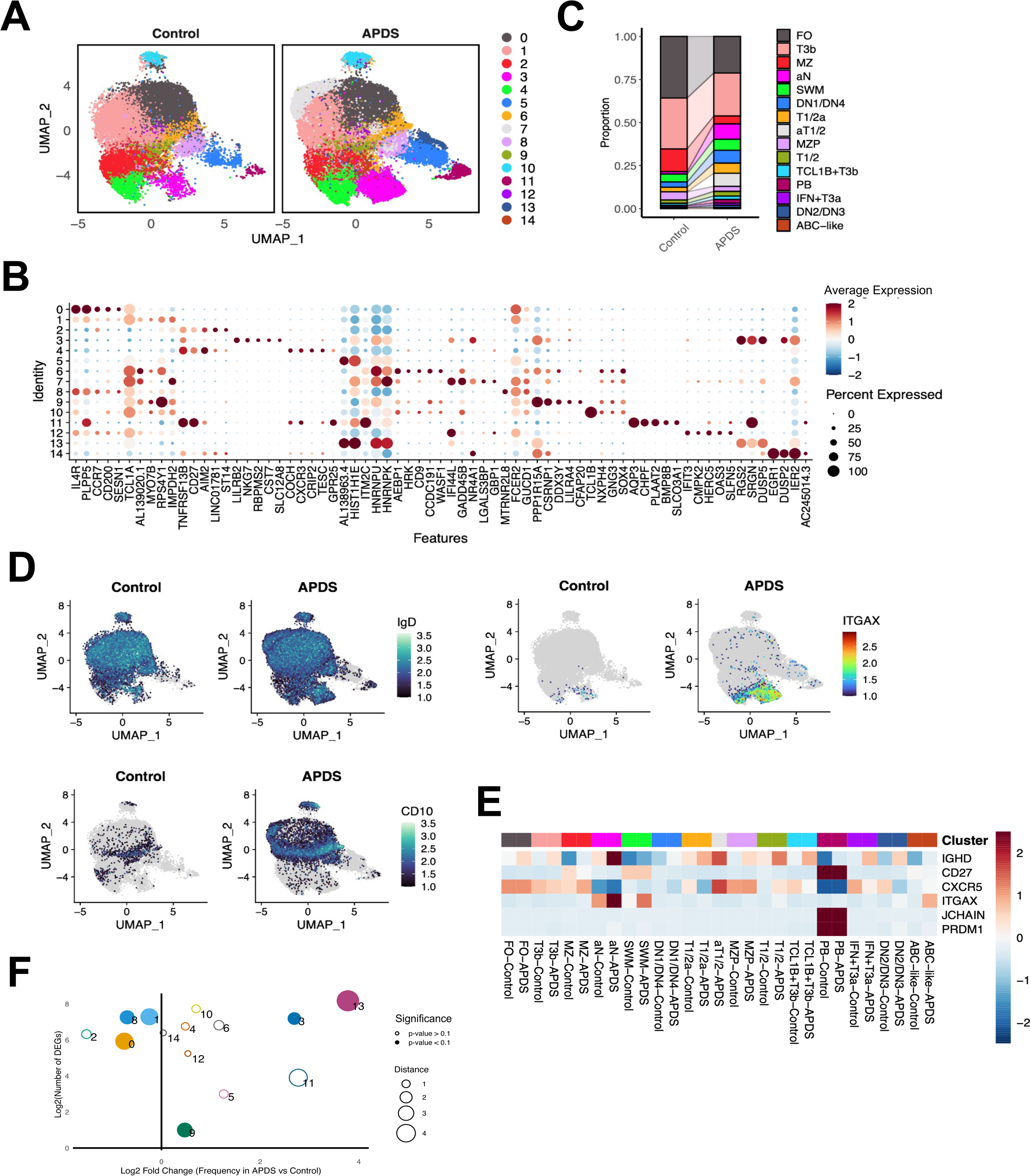
Single-cell transcriptomic and repertoire-based analysis of B cell subsets in healthy controls compared to APDS patients. CITE-seq and single-cell RNA-sequencing of the total B cell (CD3-CD19+) compartment from peripheral blood from four patients with *PI3KCD* variants causing APDS and four healthy controls. (A) UMAP projection of total B cells (CD3-CD19+) profiled by scRNA-seq annotated by control or APDS condition with 15 unique clusters (labelled 0-14). (B) Dot plot displaying differentially expressed genes through transcriptome analysis per cluster in APDS patients. Gene expression represented by dot color, while dot size indicates the percentage of cells within each cluster expressing the gene. (C) Bar graph demonstrating proportions of clusters in APDS and healthy controls. Data used to inform cluster naming as detailed in **Supplemental Table S2**. (D) UMAP projection of scCITE-seq surface markers for IgD and CD10 for healthy controls and APDS patients. UMAP projection of RNA expression of *ITGAX/CD11C* for healthy controls and APDS patients. (E) Heat map based on comparison of pseudobulk transcriptome analysis in APDS patients and healthy controls across clusters of previously published transcriptome gene expression of transitional and double negative B cells. (F) Dot plot demonstrating multi-dimensional analysis combining population frequency, transcriptional differences, and differentially expressed gene analysis between APDS patients and healthy controls. Dot size reflects transcriptional distance via PCA Euclidean distance, and filled dots indicate permutation-derived p-values < 0.1.

Given the established role of autoantibodies in the pathogenesis of autoimmune cytopenias (AIC), we next evaluated CVID patients with and without AIC. We observed an expansion of 9G4^+^ B cells between the transitional T1/2 and T3a B cell stages in patients with AIC (geometric mean 9G4 positivity 6.3 vs 15.1% in T1/2 vs T3a B cells, p=0.02). Comparing CVID patients with and without AIC, there was increased 9G4^+^ B cell frequency in CVID patients with AIC at both the T3a and T3b B cell stages (9G4 positivity 15.1 vs 5.8%, p=0.0001 at the T3a stage and 15.5 vs 6.3%, p=0.002 at the T3b stage) (**Figure 2B and Supplemental Figure 8**). An increased 9G4 frequency in CVID patients with AIC, compared to those without AIC, persisted in downstream B cell subsets that included aN, SWM, MZP, and MZ B cells.

We hypothesized that the increased frequency of 9G4^+^ transitional B cells suggests a break in tolerance at the T2 to T3a developmental transition, which may be a shared immunophenotype among different monogenic forms of CVID. We observed a significantly increased frequency of 9G4^+^ B cells in all three groups of CVID patients, including APDS, CTLA4 deficiency, and NFKB1 deficiency as compared to healthy controls. An increased frequency of 9G4^+^ B cells was first observed at the transitional T1/2 B cell stage (CTLA4 deficiency vs APDS vs NFKB1 deficiency vs healthy controls: geometric mean 4.5 vs 5.7 vs 5.3 vs 2.2%, p=0.0025), and the frequency of 9G4^+^ B cells further increased at the transitional T3a B cell stage in all three groups of CVID patients compared to healthy controls (geometric mean 7.8 vs 8.3 vs 12.6 vs 2.8%, p=0.0003) (**Supplemental Figure 9**). Overall, these data suggested that across unique and distinct categories of CVID, a common mechanism underlies the loss of B cell tolerance at a specific transitional B cell stage, driving autoimmunity in a subset of patients in each group.

### Single-cell RNA-seq analysis reveals expansion of B cell subsets in APDS

We hypothesized, based on the 9G4 flow cytometry data, that a clinically significant tolerance break occurs in transitional B cells, driving the downstream accumulation of autoreactive B cells in CVID patients with autoimmunity. We used the unbiased approach of CITE-seq and single-cell RNA-sequencing of the total B cell (CD3^-^CD19^+^) compartment from the four patients with *PI3KCD* variants causing APDS and four healthy controls.

We defined 15 clusters of B cells from APDS patients and healthy controls (labelled 0-14) (**Figure 3A**). Identification and naming of each cluster (detailed in **Supplemental Table 2**) was inferred from known differentially expressed genes as depicted in the dotplot in **Figure 3B**, and by comparison with the gene expression of transitional B cells [12] and of double negative B cell subsets [25]. In the dot plot, transcriptomes are shown by cluster for APDS patient and healthy control B cells, respectively (**Figure 3B and Supplemental Figure 10**). CITE-seq utilizing oligo-tagged antibodies for IgD and CD10 and RNA expression for *ITGAX/CD11C* are displayed in UMAP format for APDS patients and healthy controls (**Figure 3D**). A heat map profiles expression for genes that were of major interest: *IgD*, *CD27*, *CXCR5*, *ITGAX/CD11C*, J chain and *PRDM1* (**Figure 3E**). **Figure 3F** illustrates a multi-dimensional analysis combining population frequency, transcriptional differences, and differentially expressed gene analysis between APDS patients and healthy controls. This global analysis demonstrates that clusters 3 (aN) and 13 (DN2/DN3) have the most striking differences between APDS and healthy controls. Although clustering by flow cytometry for surface proteins and by transcriptome are never congruent, we used IgD, CD27, CXCR5 and CD11c since they are the four cell surface markers utilized for the initial categorization of human B cells and double negative B cells as initially described by Sanz and colleagues [26]. J chain is expressed at the highest levels in plasmablasts and plasma cells, but it is also expressed in all activated B cells, especially DN3 B cells [25].

PRDM1 (also known as BLIMP1) is the canonical transcription factor typically described in plasmablasts and plasma cells. UMAPs of additionally instructive genes of interest are shown (**Supplemental Figure 11**). These included *IGHM*, *TBX21/TBET* (a transcriptional inducer of *ITGAX*), *CD95/FAS*, which broadly represent B cells of germinal center origin [27], *TCL1A*, which in our hands marks naïve and non-germinal center derived B cells [28], *TCL1B*, which is uniquely found in transitional B cell origin cluster 10 and helps demarcate it, and *PLD4*. TCL1A and 1B are differentially expressed co-activators for AKT. *PLD4* or phospholipase D4 encodes an endosomal exonuclease that breaks down TLR9 ligands and is a good measure of relatively recent B cell activation.

As can be seen visually in **Figure 3A and Supplemental Figure 12**, Cluster 7 (aT1/2) is uniquely present in APDS and clusters 3 (aN), 4 (SWM), 5 (DN1/DN4), 6 (T1/2a), 9 (T1/2), 10 (TCL1B^+^T3b), 11 (PB), 12 (IFN^+^T3a) and 13 (DN2/DN3) are all significantly expanded in APDS subjects compared to controls (**Supplemental Table 2**).

### APDS B cells show expansion of autoreactive VH4-34^+^ transitional and extrafollicular double negative B cells

Given the expansion of activated transitional B cell subsets in APDS, we questioned whether clusters 6, 7 (a cluster not seen in controls), and 10 represented subsets of self-reactive transitional B cells that have expanded in disease state. We noted that from the T1/2 stage onwards there was a relative enrichment of VH4-34^+^ B cells in all the transitional stages including activated transitional B cells in clusters 9, 6, 7, 12 and 1 (**Figure 4A**). In APDS patients, these B cells had a higher frequency of *PLD4* positivity, suggesting relatively recent B cell activation (**Supplemental Figure 11F**). Overall, consistent with the generation of mainly extrafollicular B cell responses in APDS, a decreased frequency of somatic hypermutation was clearly observed in APDS patients, especially in clusters 3, 4, 11, and 14 (**Figure 4B**). Consistent with this finding was the relative persistence of IgM expression in APDS patient B cells across many subsets that were class-switched or IgM low in healthy controls, including the BLIMP1 positive plasmablasts (**Supplemental Figure 11A**).

**Fig 4.**
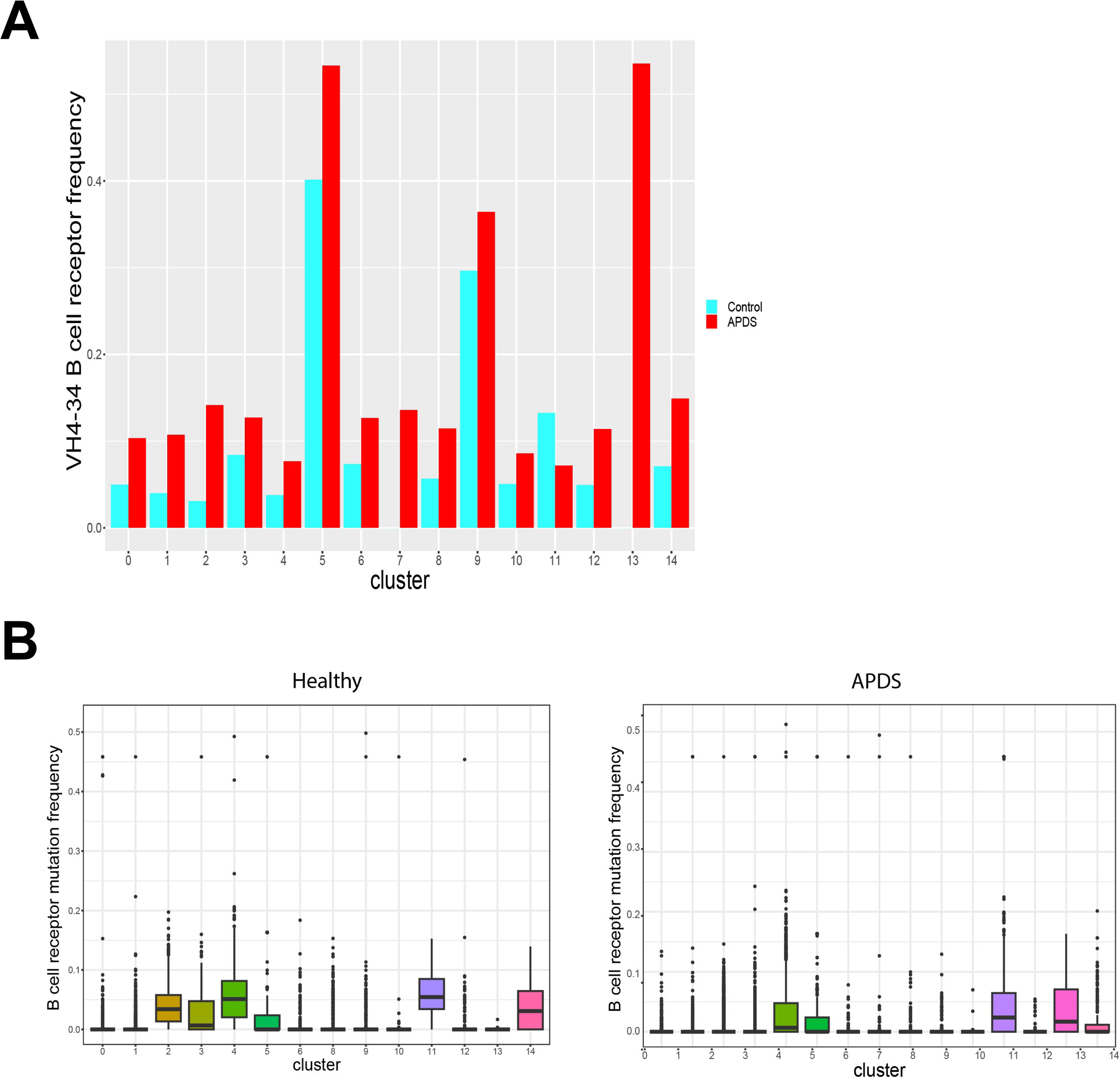
VH4-34^+^ B cell frequency is increased in APDS patients with decreased frequency of somatic hypermutation. (A) Column chart showing VH4-34 expression through B cell receptor repertoire analysis displayed by cluster for healthy controls and APDS patients. (B) Box plots showing frequency of somatic hypermutation through B cell receptor repertoire analysis by clusters for healthy controls and APDS patients.

As noted in other autoimmune diseases, in APDS there was a marked expansion of extrafollicular activated B cell subsets. For example, APDS patients had an expansion of activated Naïve B cells, cluster 3, and of double negative (DN) B cells (IgD^-^CD27^lo^), in clusters 5 and 13 (**Figure 3A**). Cluster 5 DN B cells expanded in APDS patients. Most of the cells in cluster 5 in healthy controls were *CXCR5^+^CD11c^-^* cells, and these cells have been characterized by our group and others as DN1 B cells [25, 29]. Recent evidence suggests that these DN1 B cells may represent fairly long-lived CD27^lo^ memory B cells [30]. Some B cells in cluster 5 from healthy controls were *CXCR5^+^CD11c^+^*, broadly corresponding to DN4 B cells that we have recently described [25]. Cluster 13 cells are not seen in controls but emerge in APDS patients.

The use of surface markers to categorize B cells is in principle biased, and single-cell RNA-sequencing allows unbiased categorization of these DN B cells. DN B cells were re-clustered, and we transcriptionally describe these clusters as DN.0, DN.1, DN.2 and DN.3 B cells (please note that in the single-cell data, we use the DN.n nomenclature to distinguish these subsets from those characterized by flow cytometry and cell sorting). DN.0 and DN.2 B cells are seen in healthy controls (**Supplemental Figure 13A**). DN.2 B cells expand strikingly in APDS patients and DN.1 and DN.3 B cells are only seen in APDS. While there is some increase in *ITGAX^+^* DN B cells in APDS, the most striking change in disease is the induction in disease of DN B cells that co-express *JCHAIN*, *MZB1* and *TXNDC5* (**Supplemental Figure 13B**). *TXNDC5* encodes a member of the protein disulfide isomerase family and with *MZB1* and *JCHAIN* represents genes characteristic of plasmablast-like cellular differentiation. These cells are most similar to the DN3 B cells that we have characterized by flow cytometry and transcriptionally (using bulk RNA-sequencing), and closely resemble plasmablasts in some aspects; we have shown previously that DN3 B cells accumulate in disease tissues [25].

### Clonal expansion observed across transitional and extrafollicular DN-like B cells in APDS

Relatively large B cell clonal expansions were noted in APDS subjects (**Figure 5**), and when examining the six largest clones, three were distributed across transitional and activated transitional B cell populations as well as activated Naïve B cells. Three clones in APDS patients, clones 2, 4 and 6 accumulated specifically in clusters 5 and 6 overlapping with DN.2 and DN.3 B cells that express high levels of *JCHAIN, MZB1* and *TXNDC5*. Interestingly, clusters 5 and 13 had the highest proportion of VH4-34^+^ cells, and it appears likely that clonal expansions of autoreactive B cells accumulate in the DN3-like, transcriptionally defined DN.2 and DN.3 B cell clusters.

**Fig 5.**
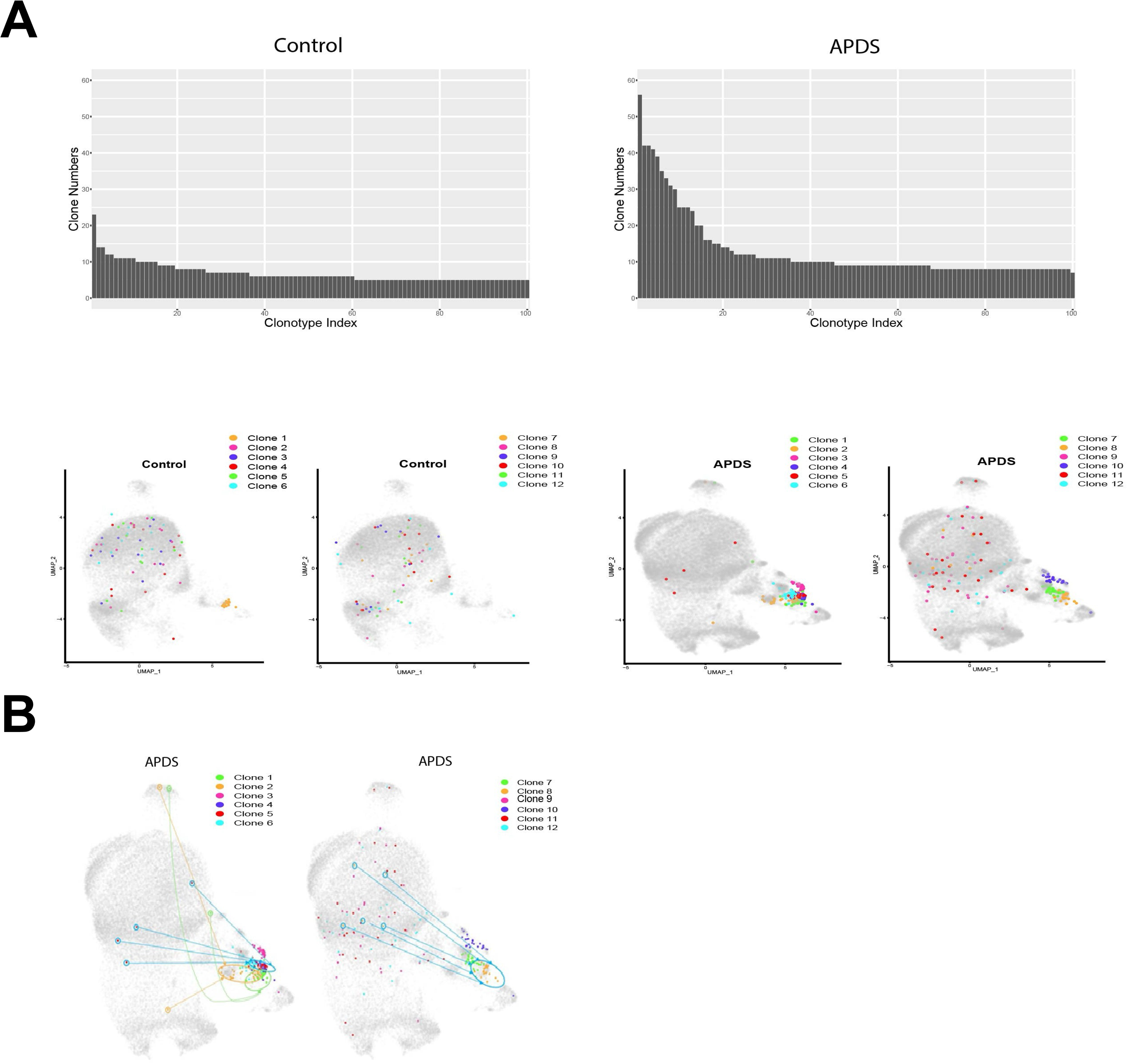
Single-cell transcriptomic analysis of double negative B cells. (A) UMAP projections of double negative (DN) DN.0, DN.1, DN.2, and DN.3 B cell clusters by single-cell RNA sequencing and (B) UMAP projections of single-cell RNA sequencing transcriptome differential gene expression of defined DN genes in healthy controls and APDS patients, including *CXCR5*, *JChain*, *ITGAX/CD11C*, *TXNDC5*, and *MZB1*. Each purple dot represents cell expression of the particular gene.

Clonal expansion was additionally observed in cluster 3, and notably there was a marked increase in B cells that co-express *ITGAX (CD11C), TBX21 (TBET)* and *FAS/CD95* in APDS patients (**Supplemental Figure 11**). These *ITGAX^+^* populations include activated Naïve B cells (cluster 3), some DN2-like B cells (cluster 13, in the DN.3 population) and CD27^+^ ABC-like cells (cluster 14). CD27^+^ ABC-like cells are defined in this report as *IgD*^-^*CD2*7^+^*CXCR5*^-^*CD11c*^+^ B cells (to distinguish them from DN2 B cells that are CD27^lo^) and these CD27^+^ B cells are generally more highly somatically mutated and are germinal center derived [31]. While a large expansion of aN B cells and DN B cells was noted in APDS, in contrast ABCs were not expanded in APDS. This supports the notion that most B cell expansion in this disease context is extrafollicular.

When examining the 12 most expanded clones in APDS patients, it became apparent that some clones that accumulate in the extrafollicular DN B cell compartment in cluster 5 in APDS patients were also observed scattered within the transitional B cell compartments, in which tolerance was initially broken. This was particularly apparent in cluster 9 (**Figure 5B**), and clonal connectivity could be established between activated B cells in the transitional B cell compartments and their related clones that accumulated in the DN B cells clusters in APDS patients (**Figure 5C**).

Finally, APDS patients develop dense lympho-infiltrative disease and thus we hypothesized that autoreactive, extrafollicularly derived B cells may infiltrate end-organs. Iterative immunofluorescence analyses of lung tissue from four patients with APDS demonstrated expansion of double negative B cells in three, consistent with our recent demonstration that the activated B cells that most readily accumulate in disease tissues are double negative B cells, especially DN3 B cells [25] (**Supplemental Figure 14**).

### High oxidative phosphorylation in unique expanded activated transitional B cells in APDS patients

While a list of all B cell clusters is provided in **Supplemental Table 2**, the following changes in APDS described in ontological sequence were of interest. APDS is defined by a relative T1/2 B cell expansion by flow cytometry [32, 33] and we did find that APDS transitional T1/2 B cells (cluster 9) were expanded in APDS. More striking was a small “activated” T1/2 B cell cluster, cluster 6, of quiescent metabolic state in both healthy controls and in APDS (**Figure 6 and Supplemental Figure 12**) which was more significantly expanded in APDS patients. A distinct expanded and metabolically active T1/2 cluster, cluster 7, was unique to APDS patients. A cluster of activated T3a-like B cells, cluster 10, characterized by the expression of *TCL1B*, that encodes a specific co-activator of AKT, expanded markedly in APDS patients. Of note, the T3-like clusters, clusters 1, 10 and 12, that are relatively metabolically inactive in healthy controls all display high relative oxidative phosphorylation status in APDS patients. Similarly, the MZP like cluster, cluster 8, that is relatively quiescent in healthy controls displays high relative Oxidative Phosphorylation in APDS patients (**Figure 6**). These transcriptomic changes strongly suggest that in APDS, T3-like B cells and MZP B cells have been activated, presumably by self-antigens and/or T cells. In addition, the expansion and the sharp increase in oxidative phosphorylation of several disease-linked activated transitional B cell subsets provide an underlying mechanism for the survival of self-reactive transitional B cells and the consequent break in immunological tolerance in these APDS patients.

**Fig 6.**
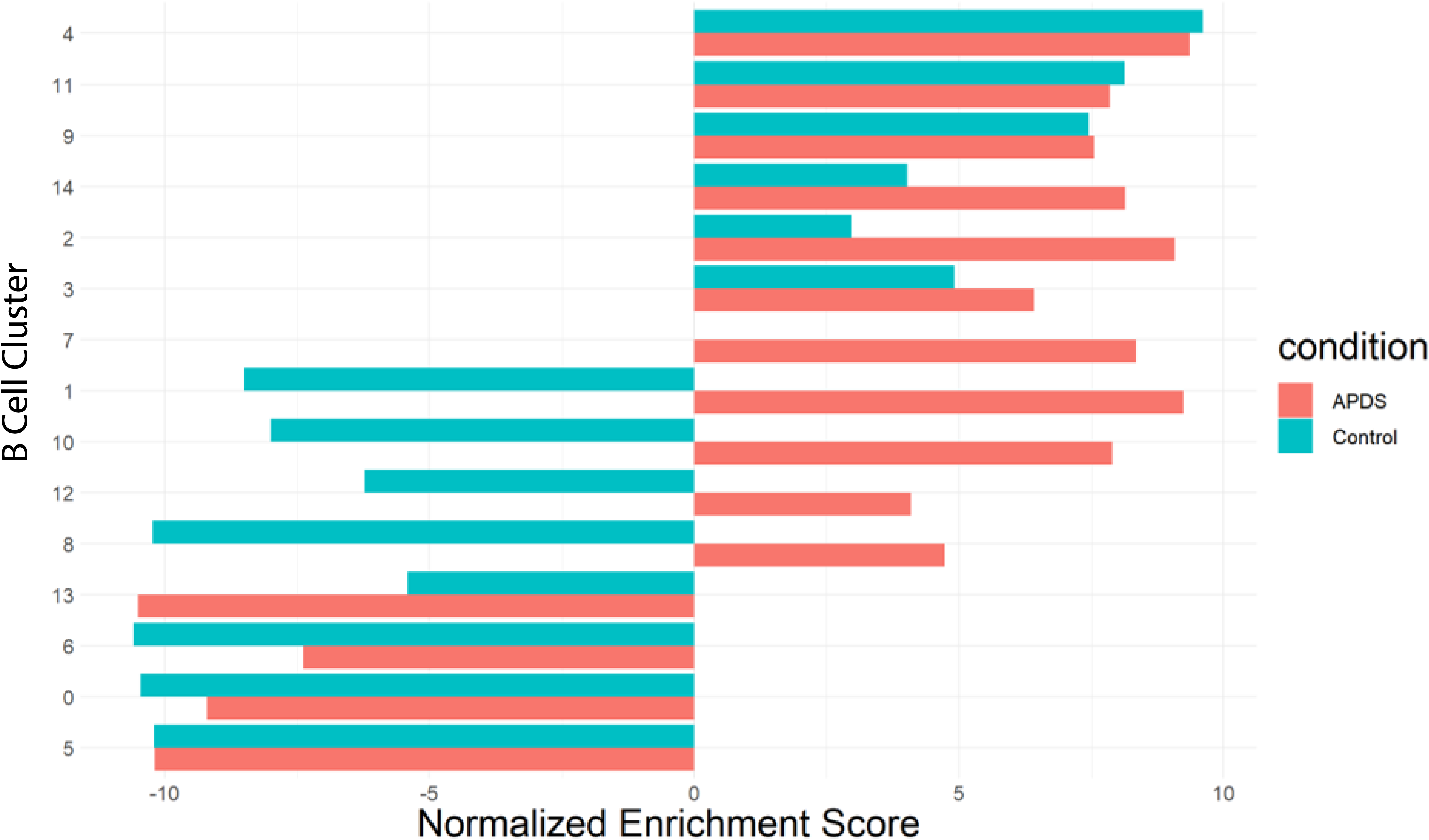
Expanded transitional B cells exhibit enhanced oxidative phosphorylation in APDS patients with autoimmunity. Bar plot displaying single-cell sequencing normalized enrichment score for the oxidative phosphorylation pathway (x axis) for each single-cell B cell cluster (y axis) for APDS patients versus healthy controls.

## DISCUSSION

The onset of autoimmunity in CVID may sometimes involve cell-intrinsic B cell changes contributing to a break in tolerance but may also be linked in other forms of the disease to the expansion of CD4^+^ T cells or the functional loss of regulatory T cells. Expanded dysregulated CD4^+^ T cells promiscuously interact with autoreactive transitional B cells that should normally be deleted or rendered quiescent by anergy. Cell intrinsic hyperactivation of B cells may also prevent the squelching or elimination of autoreactive transitional B cells. As a result, B cells normally destined for extinction or quiescence may be rescued by the induction of the transcription of anti-apoptotic genes and expand and preferentially differentiate into extrafollicular B cells. This premature and promiscuous activation of autoreactive transitional B cells may preferentially facilitate their differentiation into pathogenic extrafollicular B cells that can infiltrate end organs and contribute to tissue damage. We show here that a cell intrinsic role for NFKB1 in CD4^+^ T cells contributes to the CD4^+^ T cell dysregulation observed in patients who have inherited dysfunctional *NFKB1* variants and may contribute to autoimmunity through interaction with expanded autoreactive transitional B cells, similar to APDS [12] and CTLA4 deficiency [19].

Previously, we have shown that surface CCR7 is first upregulated on human transitional B cells at the T3a B cell stage [19]. Our data argues that expansion, activation, increase in oxidative phosphorylation, and the potential consequent anti-apoptotic protection of self-reactive transitional B cells—cells that would normally be eliminated—begins at the T2 stage. One possible anatomical explanation for this phenomenon is that, in humans, transitional T2 B cells recirculate freely, and likely have direct access to expanded CD4^+^ T cells as they enter their respective zones within the white pulp of the spleen.

Peripheral B cell tolerance is maintained through multiple checkpoints, including the induction of anergy or functional silencing, elimination of self-reactive B cells following somatic hypermutation and selection in the germinal center, and extrafollicular mechanisms involving TLR co-stimulation and dysregulated T cell help, often induced by cross-reactivity with microbial antigens [34–36]. Most PD-1^hi^CXCR5^hi^ extrafollicular helper T cells at steady state have been shown to be self-reactive [37], and these cells are normally kept in check by Tregs.

Previous studies have shown that defects in Tregs contribute to peripheral B cell tolerance breakdown. This is evident in FOXP3-deficient IPEX patients, who lack functional Tregs, as well as in patients with defects that reduce Treg numbers or impair Treg function, such as ADA deficiency or CD40L deficiency–both associated with the expansion of mature naïve autoreactive B cells [38–40]. In patients with *CTLA4* variants, loss of CTLA4 function leads to Treg dysfunction, resulting in aberrant T effector activation and inappropriate B cell stimulation by T follicular helper cells [19].

Extrafollicular T-B collaborations likely induce sufficient activation-induced cytidine deaminase (AID) expression to induce class switching and low-level somatic hypermutation. Most unswitched activated naïve B cells may be generated from the marginal zone precursor (MZP) and MZB-2 B cell pools, which can be activated in both T-independent and T-dependent pathways. In contrast, class switched DN B cells are a heterogeneous population and are thought to largely originate from activated T3 or FO B cells that have received T cell help. Activated extrafollicular B cells finally differentiate into antibody-secreting plasmablasts. We also observed a modest but significant increase in self-reactive 9G4^+^IgD^-^CD27^+^ switched memory B cells, supporting a prominent role for extrafollicular B cell responses in driving autoimmunity in CVID. However, occasional high-affinity autoantibodies may still emerge from germinal center reactions. Extrafollicular B cell activation –with expansion of activated naïve B cells and DN B cells—is well-described in SLE, and more recently has also been reported in severe COVID-19 infections [41, 42].

As immunomodulatory therapeutic options for CVID advance, there may be opportunities for therapeutic intervention in the pre-clinical disease space, as is already being explored for several autoimmune clinical entities such as rheumatoid arthritis [43]. As CVID patients who present clinically as CVID have an 11-fold higher risk of death due to auto-inflammation, a deeper understanding of the mechanisms driving immune dysregulation in these patients could inform strategies for earlier diagnosis and targeted intervention.

## MATERIALS AND METHODS

### Recruitment of patients with CVID

This research was performed in accordance with the Mass General Brigham and Beth Israel Lahey Health Institutional Review Boards and cohorts from the Massachusetts General Hospital, Beth Israel Lahey Health, the University of South Florida (protocols 2011P000940, 20243028, and 2018P001584). Written informed consent for specimen collection was obtained from all patient participants included in this study. From primary immune deficiency cohorts followed at the Massachusetts General Hospital [11] and Beth Israel Lahey Health patients underwent genotyping by next-generation panel or whole exome sequencing.

### Inclusion criteria for CVID cases

Patients were defined as CVID using the following criteria based on clinical evaluation by an immunologist with expertise in CVID:

- CVID:

Reduced, age-adjusted total serum concentrations of IgG and IgA and/or IgM
Abnormal specific antibody response to immunization
The absence of any other defined immunodeficiency state (i.e., CVID is a diagnosis of exclusion)

Patients were defined as having clinical autoimmunity by the presence of at least one of the following criteria: Autoimmune enteritis, nodular regenerative liver hyperplasia, granulomatous lymphocytic interstitial lung disease, autoimmune cytopenias (AIC), inflammatory arthritis, and/or secondary lymphoid organ hypertrophy. Chronic cytopenias, specifically, were defined as positive history on physician chart review or otherwise as captured by review of all cytology in the electronic medical record as total hemoglobin, platelet count, absolute neutrophil count, less than age-matched reference range and persistent >12 months or positive autoantibodies (direct antiglobulin test, anti-platelet and/or anti-neutrophil) as previously described [11, 44–46].

Patients with monogenic CVID included CTLA4 deficiency (n=11), APDS (n=4), and NFKB1 deficiency (n=5). Patient variants were validated as listed in **Supplemental Table 1**, row 3.

Patients were divided into different groups for various analyses, including patients with monogenic CVID with an autoimmune clinical phenotype (n=16), and patients with monogenic CVID without an autoimmune clinical phenotype (n=4). We also divided CVID patients without AIC (n=11), and with AIC (n=9). Twelve healthy controls were included.

We studied nine patients with *NFKB1* variants that encodes the NF-κB1 p105 subunit that is processed into the p50 transcription factor. Variants were identified that spanned each domain of *NFKB1* **(Supplemental Figure 1A).** The variants in *NFKB1* included a nucleotide substitution that may affect splicing (c.####-#N > N, p.G55Ifs*8), one nonsense variant (c.####-#N > N, p.Q468*), four frameshift deletions (p.G55Ifs*8, p.Asp452Glufs*5(c.1356del), p.T869Vfs*10(c.2602_2603dupGG), pGly397Alafs*35(c.1190del) (**Supplemental Table 1**).

Several of the variants were also missense variants in the p50 RHD domain (p.R231H(c.692 G>A), p.G261R(c.781G>C), p.R335Q(c.), p.D452Efs*5(c.1356del)), the p105 linker region (p.Q468X(c.), p.N515S(c.1544 A>G)), and the p105 death domain (p.T869Vfs*10(c.2602_2603dupGG)).

To determine whether the *NFKB1* variants in our CVID cohort (**Supplemental Figure 1A)** were loss-of-function variants, we expressed the variants using the pSBbi-GP vector (Addgene #60511), which uses the Sleeping Beauty transposon and a bidirectional promoter to thwart silencing of the expression of the gene of interest and integrated them into Jurkat cells using the Sleeping Beauty transposase SB100x (Addgene #34879). In this construct, the EF1alpha promoter drives expression of *NFKB1* variant, whereas the RPSBA promoter drives expression of eGFP and the puromycin resistance gene to track cells that have the integrated construct. These cells were selected with puromycin and sorted for high GFP positivity, after which they were subjected to RNA-seq. Transcriptomic profiling revealed that of the seven variants cloned and expressed, all but one were highly similar to the empty vector pSBbi-GP, whereas the transcriptional profile of R335Q was most similar to that of wild-type *NFKB1* (**Supplemental Figure 1B)**. Closer examination and searches of *NFKB1* R335Q from gnomAD and other databases showed that this *NFKB1* variant is common in the population and therefore may be an incidental variant that was identified in exome sequencing of the MGB cohort.

Comparison of the transcriptomic profiling of the loss-of-function *NFKB1* variants and examining the enriched pathways in the differentially expressed genes between the loss-of-function *NFKB1* variants and the intact/wild-type NFKB1 showed that signatures associated with ribosome biogenesis, ncRNA and rRNA metabolism and processing were downregulated upon overexpression of wild-type NFKB1 (**Supplemental Figure 1C**). Pathways associated with Myc targets, MTORC1 signaling, E2F targets, and the unfolded protein response were all downregulated in cells in which functional NFKB1 was overexpressed, whereas the expression of the loss-of-function *NFKB1* variants (and empty vector) were associated with gene sets in which hypoxia, MTORC1 signaling, and glycolysis were upregulated (**Supplemental Figure 1D**). Additionally, enriched pathways in DEGs associated with ribonucleoprotein complex biogenesis, ribosome biogenesis, RNA processing and metabolic process (including ncRNA and rRNA processing) were all downregulated upon expression of functional NFKB1. Taken together, these data suggest that *NFKB1* overexpression may serve to act as a transcriptional repressor, leading to the decrease in metabolism and translation, as corroborated by the downregulation of enriched pathways associated with Myc targets, E2F targets, MTORC1 signaling, and the unfolded protein response, but certain CVID linked loss-of function-variants fail to show this repressive activity.

### Flow cytometry

The distinction between transitional and follicular human B cells was accomplished using ABC transporter efflux profiling in combination with CD73 surface profiling as previously described [11, 12, 47, 48]. In brief, 5 million frozen peripheral blood mononuclear cells (PBMCs) were incubated in RPMI 1640 media containing MitoTracker green (MTG) (Thermo Fisher Scientific) at a 1:10,000 dilution (100-nM concentration) for 30 min at 37°C. Subsequently, cells were spun down at 1250 rpm for 6 min and rested in fresh RPMI 1640 media for an additional 30 mins at 37°C to allow efflux of the dye in ABC^+^ cells. Cells were blocked for 15 min with human Fc block (BD Biosciences) at a concentration of 2.5 µg per 1 million cells. Cells were surface-stained with optimized concentrations of fluorochrome-conjugated primary antibodies for 30 min at 4°C. To minimize the interaction of polymeric dyes and improve discrete fluorochrome readout, Brilliant Stain Buffer Plus (BD Biosciences) was included at 10 µL per condition during surface staining. DRAQ7^TM^ (BioLegend) was used to exclude dead cells as per the manufacturer’s instructions. Compensation was performed using VersaComp Antibody Capture Beads (Beckman Coulter) apart from MTG, which required staining of Ramos cells [American Type Culture Collection (ATCC)] with MTG for ≥30 min at 37°C, and 9G4 antibody required Flow Cytometry Protein G Binding Beads (Bangs Laboratories). 9G4 antibody was obtained from IGM Biosciences and conjugated to APC fluorophore using APC Mix-n-Stain Fluorescent Protein & Tandem Dye Antibody Labeling Kit using kit instruction (Biotium). For analysis, B cells were run on a five-laser BD FACS Symphony (5S Symphony). Gating was performed as shown in **Supplemental Figure 2**. Antibody-secreting cells were not included in the analysis due to comprising <1% of total B cells in over 75% of samples tested. For sorting, B cells were run on a Special Order Research Products Aria II (BD Biosciences) and collected in RMPI 1640 media supplemented with 20% fetal bovine serum (FBS). B cell panel was as follows: CD3 (UCHT1, BD Biosciences, 1:30), CD19 (SJ25C1, BioLegend, 1:30), CD27 (L128, BioLegend, 1:40), IgD (IA6-2, BioLegend, 1:80), CD38 (HIT2, BioLegend, 1:30), CD24 (ML5, BioLegend, 1:60), CD10 (HI10a, BioLegend, 1:20), CD45RB (MEM-55, Thermo Fisher Scientific, 1:20), CD21 (B-ly4, BD Biosciences, 1:20), IgM (G20-127, BD Biosciences, 1:80), IgG (G18-145, BD Biosciences, 1:100), CD20 (2H7, BD Biosciences, 1:60), CD73 (AD2, BD Biosciences, 1:40), CXCR5 (J252D4, BioLegend, 1:30), CD11c (B-ly6, BD Biosciences, 1:20), 9G4 (IGM, 1:200).

We defined our B cell populations as transitional 1/2 (T1/2: IgD^+^CD27^-^CD10^+^MTG^+^), transitional 3a (T3a: IgD^+^CD27^-^CD10^-^CD73^-^MTG^+^), transitional 3b (T3b: IgD^+^CD27^-^CD10^-^ CD73^+^MTG^+^), follicular (FO: IgD^+^CD27^-^CD10^-^CD73^+^MTG^-^), activated naïve (IgD^+^CD27^-^ CD10^-^CD45RB^-^CD21^lo^CD11c^hi^), MZP (IgD^+^CD27^-^CD10^-^CD45RB^+^), marginal zone (IgD^+^CD27^+^), switched memory (IgD^-^CD27^+^), and double negative (IgD^-^CD27^-^) as previously defined by flow cytometry.[12]

### CITE-Seq (Cellular Indexing of Transcriptomes and Epitopes by Sequencing) and 10X single-cell sequencing

CITE-seq was performed according to 10X protocol as previously described [49]. Live B cells (CD19^+^CD3^-^7AAD^-^) were sorted prior to cell staining. B cell fluorophore panel was as follows: CD3 (UCHT1, 1:50), CD19 (SJ25C1, 1:30). After sorting, total live B cells were incubated with Human Fc Block (BD Biosciences) for 10 minutes on ice. They were then stained with TotalSeq-C antibodies (BioLegend) for 30 minutes on ice along with hashing antibodies (BioLegend). Total-Seq antibody panel was as follows: CD73 (AD2; 1:200), IgD (1A6-2, 1:400), CD45RB (MEM-55, 1:100), CD10 (HI10a, 1:100). The single cell suspensions had 72-100% viability and was diluted at a concentration of 10 million cells/uL. All eight suspensions were pooled together and then split into four technical replicates to reduce batch effects. Cells were processed using the Chromium Single Cell 5’kit v2 and single cell emulsions were generated. The antibody-derived tagged libraries were created as detailed in the TotalSeq-C Antibodies Hashing with 10X Single Cell 5’ Kit v2 protocol (BioLegend).

### Single-cell RNA-seq Data Processing

scRNA-seq data and CITE-seq data were processed in R (v4.1.2)[50] using the Seurat package (v4.1.0).[51–53] Raw UMI count matrices generated from the Cellranger 10X pipeline were loaded and merged into a single Seurat object after demultiplexing all samples using hashing information from the CITE-seq antibodies.

We filtered cells based on the following parameters: cells with less than 1000 RNA transcripts detected, with less than 250 gene counts, with a 20% or greater fraction of total gene counts mitochondrial, or with a greater than 80% ratio of log10 gene counts and RNA transcript counts.

Genes expressed in fewer than 10 cells, AC233755.1 (IGHV4-38-2), AC233755.2 (IGHV3-43D) and immunoglobulin genes except 9G4 (IGHV4-34), IGHA1, IGHA2, IGHG1, IGHG2, IGHG3, IGHG4, IGHE, IGHD, and IGHM were extirpated from the gene counts matrix. After scaling and normalizing the gene counts, a PCA plot based on the 3000 most variable genes found mitochondrial genes significantly affecting the PCA plot, but not the cell cycle genes. Therefore, the collective object was split into 8 objects, one for each sample, and SCTransform was applied to each with the regression of mitochondrial genes to mitigate their effects subsequent analysis. Seurat’s SCTransform mitigates technical noise in the count data with pearson residuals generated from a unique regularized negative binomial model to mitigate overfitting. [54]

We took the most variable SCTransformed genes shared by all samples to generate PCA PCs and used 40 of the PCs to generate uniform manifold approximation and projections (UMAPs). For clustering, we calculated the k nearest neighbors shared nearest neighbor (SNN) and used the Louvain algorithm, an unsupervised modularity optimization-based clustering algorithm for community finding. After several tests of different resolution values, we decided upon the resolution of 0.5. Two small clusters that contained markers for T cells and macrophages were removed, and the PCA, UMAP, and clustering was redone in the same manner.

To assess global changes in the dataset between APDS disease and healthy controls, we performed a Principal Component Analysis (PCA) on pseudobulk counts, accounting for Seurat clusters and disease state by individual patients. Euclidean distances between patients were calculated based on the first two principal components (PC1 and PC2), measuring divergence between APDS and Control groups in PCA space. To determine the statistical significance of the observed distances, we conducted permutation testing if the differences they observed between the disease and healthy control groups were statistically significant, rather than occurring by chance. Empirical p-values were computed as the proportion of permuted distances that were greater than or equal to the observed distance.

For differential gene analysis, we ran FindAllMarkers to find the top most differentiated gene expressions for each cluster. The analysis was restricted to genes expressed in at least 20% of cells in either cluster and with a minimum difference in detection rate of 10%. Only genes with an adjusted p-value less than 0.01 were considered significant. Top markers were used to generate a heatmap.

For Gene Enrichment Analysis, we further divided each cluster by their Control or APDS condition cells and ran a differential gene analysis. We used the Molecular Signatures Database (MSigDB) from *Gene Set Enrichment Analysis* (GSEA) to identify differential enrichment of Hallmark cellular and metabolic pathways between the clusters of different conditions.

For clonality analysis, we used scRepertoire (v1.8.0) for the 10XGenomics BCRannotated files and merged it with the single-cell data with combineBCR.[55] As DN B cells have undergone hypermutation to a limited degree, to trace their potential non mutated precursor B cells, we defined BCR clonotypes by their heavy and light chain VDJC allele combination. For somatic hypermutation analysis, we processed the 10X BCR annotations through IMGT HighV-QUEST and used the R package shazam (v1.1.2), a component of the Immucantation analysis framework.[56]

### Definitions

We identified patient-enriched B cell clusters defined as clusters with a ratio of frequency of cells per cluster compared to total cells of patients to controls of >1.25.

We analyzed immunoglobulin isotypes based on immunoglobulin heavy chain transcripts in total B cells. We identified class-switching based on %IgM, IgA and IgA in APDS and controls; if <85% IgM, considered unswitched.

We analyzed BCR expression for clonal expansion, defined as a clonal threshold of >4 unique BCR sequences. In healthy controls, clusters were defined as extrafollicular-derived (EF-derived) if the median somatic hypermutation (SHM) was zero, and germinal center-derived (GC-derived) if the median SHM was greater than zero. In APDS, clusters were defined as EF-derived if the SHM was significantly lower than healthy SHM, and GC-derived for clusters with median >0 and not significant compared to healthy controls (**Supplemental Table 2**).

### Microscopy

#### Multi-color immunofluorescence staining

For multi-color immunofluorescence staining, tissue samples were fixed in formalin, embedded in paraffin, and 4 μm sectioned. The tissue sections were deparaffinized in xylene and rehydrated by serial passage through graded concentrations of ethanol. Endogenous peroxidase in tissues was blocked with 0.3% H2O2/methanol for 10 min. Heat-induced epitope retrieval was performed for 5 min at 95 °C in Tris–EDTA pH 9.0 buffer or Citrate pH 6.0 buffer. Sections were washed in cold running tap water and Tris-buffered saline–0.05% Tween20 (TBST) and blocked with blocking/antibody diluent for 8 min. Specimens were incubated with primary antibodies specific for the following proteins: anti-CD19 (1:200; SKU310; Biocare Medical), anti-IgD (1:2000; AA093; DAKO), anti-CD27 (1:1000; clone: ab131254; Sigma Prestige), anti-CD73 (1:200; clone: D7F9A; Cell Signaling Technology). Then, sections were incubated with polymer HRP Mouse + Rabbit for 10 min, followed by incubation with an Opal fluorophore from Opal™ Multiplex Kit (Perkin Elmer) for 10 min. Bound primary and secondary antibodies were then eluted with heat-induced epitope retrieval treatment with stripping buffer. After washing in cold running tap water and TBST, the process of staining and antibody removal was repeated using a different Opal fluorophore. Finally, the samples were mounted with ProLong Diamond Antifade mountant after DAPI staining (Invitrogen).

### Microscopy and Quantitative Image Analysis

Images of the tissue specimens were acquired using the TissueFAXS platform (TissueGnostics). For quantitative analysis, the entire area of the tissue was acquired as a digital grayscale image in five channels with filter settings for FITC, Cy3, Cy5 and AF750 in addition to DAPI. Cells of a given phenotype were identified and quantitated using the TissueQuest software (TissueGnostics), with cut-off values determined relative to the positive controls. This microscopy-based multicolor tissue cytometry software permits multicolor analysis of single cells within tissue sections similar to flow cytometry. Although the software has been developed and validated more recently, the principle of the method and the algorithms used have been described in detail elsewhere [57].

### Statistical Analysis

Descriptive statistics including mean, standard deviation, geometric mean, 95% confidence interval, median, interquartile range (IQR), frequency, and percentage were used to summarize the data. The Mann-Whitney test was used to compare outcomes between two groups. For comparisons involving more than two groups, ANOVA or Kruskal-Wallis tests with appropriate post hoc analyses were applied, depending on whether the data followed a normal or non-normal distribution. All p-values were two-sided, and p-values less than 0.05 were considered statistically significant. All statistical analyses were performed using GraphPad Prism, version 9.

## Supplemental Materials

**Fig. S1:** Identification of pathogenic variants of NFKB1 in an *in vitro* system allows for the identification of pathogenic *NFKB1* VUS.

**Fig. S2:** Gating strategy for flow cytometry of human peripheral B cell subsets.

**Fig. S3:** Accumulation of transitional B cells in CVID patients with pathogenic *NFKB1* variants.

**Fig. S4:** Knockout of NFKB1 in primary CD4^+^ T cells causes increased proliferation and expansion.

**Fig. S5:** Patients with pathogenic *NFKB1* variants exhibit a relative increase in effector/memory CD4^+^ T cells and a relative decline in regulatory T cells.

**Fig. S6:** Frequency of B cell subsets in peripheral blood in CVID patients with and without autoimmunity. Flow cytometric analysis of peripheral blood B cells populations in CVID patients with (n=16) and without (n=4) autoimmunity, compared to healthy controls (n=12).

**Fig. S7:** 9G4^+^ B cell frequency as percentage of CD19^+^ total B cells in CVID patients with and without autoimmunity and controls.

**Fig. S8:** 9G4^+^ B cell frequency as percentage of CD19^+^ total B cells in CVID patients with and without autoimmune cytopenias (AIC) and controls.

**Fig S9:** 9G4^+^ B cell frequency as percentage of CD19^+^ total B cells in CVID patients, subset by monogenic disease type.

**Fig S10:** Transcriptome heatmap of differentially expressed genes by cluster for healthy controls. Heatmap of scaled gene expression of the top differentially expressed genes between annotated clusters in healthy controls.

**Fig S11:** UMAP projection of **s**ingle-cell transcriptomic analysis of selected gene expression for healthy controls and APDS patients.

**Fig S12**: (A) UMAP projection of B cell clusters by single-cell RNA sequencing for individual patients with APDS (n=4) and healthy controls (n=4). (B) Proportion of cells in cluster by sample group (APDS versus healthy controls) and significance performed by Fisher’s exact test.

**Fig S13:** UMAP projection of single-cell transcriptomic analysis of selected gene expression for healthy controls and APDS patients in the DN B cell compartment.

**Fig S14:** Expansion of double negative B cells in lung tissue in APDS patients. (A) Quantitation of B cells by percentage of total B cells per sample in infiltrating lung tissue in APDS patients (n=4), showing the relative distribution of IgD^-^CD27^-^ (double negative), IgD^-^CD27^+^ (switched memory), IgD^+^CD27^+^ (marginal zone), and IgD^+^CD27^-^ (naive). (B) Representative multi-color immunofluorescence images of CD19 (red), CD27 (green), IgD (purple) and DAPI (blue) staining in lung tissue from APDS patients (n=4).

**Table S1:** Clinical characteristics of patients with monogenic CVID.

**Table S2:** List of B cell cluster designations as annotated by single-cell CITE-seq surface marker expression, single-cell RNA-sequencing differential gene expression, and gene set enrichment analysis for oxidative phosphorylation pathway. Final column designates if cluster is expanded in APDS patients compared to healthy controls.

## Acknowledgments

We thank the tissue processing core at the Ragon Institute for the processing and biobanking of the patient samples. **Funding** was provided by American Society of Hematology Research Training Award for Fellows, NIH 5T32HL007574-38, and Sala Elbaum Pediatric Research Scholars Program (KH), Howard Hughes Medical Institute Gilliam Fellowship (KZ), K23AI163350 (SB), NIH U19 AI110495 and the Ragon Insitute (SP), NIH/NAIDS P30-AI 060354 (MG) and NIH T32-HL116275, and the American Academy of Allergy, Asthma, & Immunology (AAAAI) faculty development award and an investigator-initiated research grant from Pharming (JRF).

## Author contributions

Conceptualization: KH, GY, JRF, SP Patient Sample Contribution: JRF, SB, JW Methodology: JRF, KH, GY, KZ, HAC, AH Funding acquisition: KH, KZ, JRF, SP Supervision: JRF, SP Writing – original draft: KH, GY, JRF, AH Writing – review & editing: all authors

## Data and materials availability

All data needed to evaluate the conclusions in the paper are present in the main paper or the supplemental materials.

## References

1. Woodruff, M.C., et al., Dysregulated naive B cells and de novo autoreactivity in severe COVID-19. Nature, 2022. 611(7934): p. 139–147.

2. Burnett, D.L., et al., Clonal redemption and clonal anergy as mechanisms to balance B cell tolerance and immunity. Immunol Rev, 2019. 292(1): p. 61–75.

3. Lanz, T.V., et al., Clonally expanded B cells in multiple sclerosis bind EBV EBNA1 and GlialCAM. Nature, 2022. 603(7900): p. 321–327.

4. Bonilla, F.A., et al., International Consensus Document (ICON): Common Variable Immunodeficiency Disorders. J Allergy Clin Immunol Pract, 2016. 4(1): p. 38–59.

5. Resnick, E.S., et al., Morbidity and mortality in common variable immune deficiency over 4 decades. Blood, 2012. 119(7): p. 1650–7.

6. Chapel, H., et al., Common variable immunodeficiency disorders: division into distinct clinical phenotypes. Blood, 2008. 112(2): p. 277–86.

7. Resnick, E.S. and C. Cunningham-Rundles, The many faces of the clinical picture of common variable immune deficiency. Curr Opin Allergy Clin Immunol, 2012. 12(6): p. 595–601.

8. Gathmann, B., et al., Clinical picture and treatment of 2212 patients with common variable immunodeficiency. J Allergy Clin Immunol, 2014. 134(1): p. 116–26.

9. Fischer, A., et al., Autoimmune and inflammatory manifestations occur frequently in patients with primary immunodeficiencies. J Allergy Clin Immunol, 2017. 140(5): p. 1388–1393 e8.

10. Boileau, J., et al., Autoimmunity in common variable immunodeficiency: correlation with lymphocyte phenotype in the French DEFI study. J Autoimmun, 2011. 36(1): p. 25–32.

11. Farmer, J.R., et al., Common Variable Immunodeficiency Non-Infectious Disease Endotypes Redefined Using Unbiased Network Clustering in Large Electronic Datasets. Front Immunol, 2017. 8: p. 1740.

12. Farmer, J.R., et al., Induction of metabolic quiescence defines the transitional to follicular B cell switch. Sci Signal, 2019. 12(604).

13. Berbers, R.M., et al., Chronically Activated T-cells Retain Their Inflammatory Properties in Common Variable Immunodeficiency. J Clin Immunol, 2021. 41(7): p. 1621–1632.

14. Lui, V.G., et al., Dysregulated Lymphocyte Antigen Receptor Signaling in Common Variable Immunodeficiency with Granulomatous Lymphocytic Interstitial Lung Disease. J Clin Immunol, 2023.

15. Fraz, M.S.A., et al., Raised Serum Markers of T Cell Activation and Exhaustion in Granulomatous-Lymphocytic Interstitial Lung Disease in Common Variable Immunodeficiency. J Clin Immunol, 2022. 42(7): p. 1553–1563.

16. Klocperk, A., et al., Distinct CD8 T Cell Populations with Differential Exhaustion Profiles Associate with Secondary Complications in Common Variable Immunodeficiency. J Clin Immunol, 2022. 42(6): p. 1254–1269.

17. Gereige, J.D. and P.J. Maglione, Current Understanding and Recent Developments in Common Variable Immunodeficiency Associated Autoimmunity. Front Immunol, 2019. 10: p. 2753.

18. Thauland, T.J., et al., Case Study: Mechanism for Increased Follicular Helper T Cell Development in Activated PI3K Delta Syndrome. Front Immunol, 2019. 10: p. 753.

19. Allard-Chamard, H., et al., Congenital T cell activation impairs transitional to follicular B cell maturation in humans. Blood Adv, 2024.

20. Allard-Chamard, H., et al., Congenital T cell activation impairs transitional to follicular B cell maturation in humans. bioRxiv, 2024: p. 2024.02.08.579495.

21. Pugh-Bernard, A.E., et al., Regulation of inherently autoreactive VH4-34 B cells in the maintenance of human B cell tolerance. J Clin Invest, 2001. 108(7): p. 1061–70.

22. Cappione, A, 3rd., et al., Germinal center exclusion of autoreactive B cells is defective in human systemic lupus erythematosus. J Clin Invest, 2005. 115(11): p. 3205–16.

23. Lau, A., et al., Activated PI3Kdelta breaches multiple B cell tolerance checkpoints and causes autoantibody production. J Exp Med, 2020. 217(2).

24. Richardson, C.T., et al., Failure of B Cell Tolerance in CVID. Front Immunol, 2019. 10: p. 2881.

25. Allard-Chamard, H., et al., Extrafollicular IgD(-)CD27(-)CXCR5(-)CD11c(-) DN3 B cells infiltrate inflamed tissues in autoimmune fibrosis and in severe COVID-19. Cell Rep, 2023. 42(6): p. 112630.

26. Kaminski, D.A., et al., Advances in human B cell phenotypic profiling. Front Immunol, 2012. 3: p. 302.

27. Hao, Z., et al., Fas receptor expression in germinal-center B cells is essential for T and B lymphocyte homeostasis. Immunity, 2008. 29(4): p. 615–27.

28. Narducci, M.G., et al., Regulation of TCL1 expression in B- and T-cell lymphomas and reactive lymphoid tissues. Cancer Res, 2000. 60(8): p. 2095–100.

29. Jenks, S.A., et al., Distinct Effector B Cells Induced by Unregulated Toll-like Receptor 7 Contribute to Pathogenic Responses in Systemic Lupus Erythematosus. Immunity, 2018. 49(4): p. 725–739 e6.

30. Zurbuchen, Y., et al., Human memory B cells show plasticity and adopt multiple fates upon recall response to SARS-CoV-2. Nat Immunol, 2023. 24(6): p. 955–965.

31. Lau, D., et al., Low CD21 expression defines a population of recent germinal center graduates primed for plasma cell differentiation. Sci Immunol, 2017. 2(7).

32. Lucas, C.L., et al., Dominant-activating germline mutations in the gene encoding the PI(3)K catalytic subunit p110delta result in T cell senescence and human immunodeficiency. Nat Immunol, 2014. 15(1): p. 88–97.

33. Coulter, T.I., et al., Clinical spectrum and features of activated phosphoinositide 3-kinase delta syndrome: A large patient cohort study. J Allergy Clin Immunol, 2017. 139(2): p. 597–606 e4.

34. Nemazee, D., Mechanisms of central tolerance for B cells. Nat Rev Immunol, 2017. 17(5): p. 281–294.

35. Brink, R. and T.G. Phan, Self-Reactive B Cells in the Germinal Center Reaction. Annu Rev Immunol, 2018. 36: p. 339–357.

36. Shlomchik, M.J., Sites and stages of autoreactive B cell activation and regulation. Immunity, 2008. 28(1): p. 18–28.

37. Lee, V., et al., The endogenous repertoire harbors self-reactive CD4(+) T cell clones that adopt a follicular helper T cell-like phenotype at steady state. Nat Immunol, 2023. 24(3): p. 487–500.

38. Kinnunen, T., et al., Accumulation of peripheral autoreactive B cells in the absence of functional human regulatory T cells. Blood, 2013. 121(9): p. 1595–603.

39. Sauer, A.V., et al., Defective B cell tolerance in adenosine deaminase deficiency is corrected by gene therapy. J Clin Invest, 2012. 122(6): p. 2141–52.

40. Herve, M., et al., CD40 ligand and MHC class II expression are essential for human peripheral B cell tolerance. J Exp Med, 2007. 204(7): p. 1583–93.

41. Jenks, S.A., et al., Extrafollicular responses in humans and SLE. Immunol Rev, 2019. 288(1): p. 136–148.

42. Woodruff, M.C., et al., Extrafollicular B cell responses correlate with neutralizing antibodies and morbidity in COVID-19. Nat Immunol, 2020.

43. Greenblatt, H.K., et al., Preclinical rheumatoid arthritis and rheumatoid arthritis prevention. Curr Opin Rheumatol, 2020. 32(3): p. 289–296.

44. Park, S.H., Diagnosis and treatment of autoimmune hemolytic anemia: classic approach and recent advances. Blood Res, 2016. 51(2): p. 69–71.

45. Lambert, M.P. and T.B. Gernsheimer, Clinical updates in adult immune thrombocytopenia. Blood, 2017. 129(21): p. 2829–2835.

46. Youinou, P., et al., Diagnostic criteria for autoimmune neutropenia. Autoimmun Rev, 2014. 13(4-5): p. 574–6.

47. Palanichamy, A., et al., Novel human transitional B cell populations revealed by B cell depletion therapy. J Immunol, 2009. 182(10): p. 5982–93.

48. Wirths, S. and A. Lanzavecchia, ABCB1 transporter discriminates human resting naive B cells from cycling transitional and memory B cells. Eur J Immunol, 2005. 35(12): p. 3433–41.

49. Stoeckius, M., et al., Simultaneous epitope and transcriptome measurement in single cells. Nature Methods, 2017. 14(9): p. 865-+.

50. R: A Language and Environment for Statistical Computing. 2020, R Foundation for Statistical Computing.

51. Hao, Y., et al., Integrated analysis of multimodal single-cell data. Cell, 2021. 184(13): p. 3573–3587 e29.

52. Stuart, T., et al., Comprehensive Integration of Single-Cell Data. Cell, 2019. 177(7): p. 1888–1902 e21.

53. Butler, A., et al., Integrating single-cell transcriptomic data across different conditions, technologies, and species. Nat Biotechnol, 2018. 36(5): p. 411–420.

54. Hafemeister, C. and R. Satija, Normalization and variance stabilization of single-cell RNA-seq data using regularized negative binomial regression. Genome Biol, 2019. 20(1): p. 296.

55. Borcherding, N., N.L. Bormann, and G. Kraus, scRepertoire: An R-based toolkit for single-cell immune receptor analysis. F1000Res, 2020. 9: p. 47.

56. Gupta, N.T., et al., Change-O: a toolkit for analyzing large-scale B cell immunoglobulin repertoire sequencing data. Bioinformatics, 2015. 31(20): p. 3356–8.

57. Ecker, R.C. and G.E. Steiner, Microscopy-based multicolor tissue cytometry at the single-cell level. Cytometry A, 2004. 59(2): p. 182–90.

58. Allard-Chamard H et al. Congenital T cell activation impairs transitional to follicular B cell maturation in humans. BIORXIV, 2024/579495_

