## Supplemental Materials for "Loss of B cell tolerance at the T2/T3a B cell transition is a convergent pathogenic mechanism in common variable immunodeficiency"

### Supplemental Figure 1

A

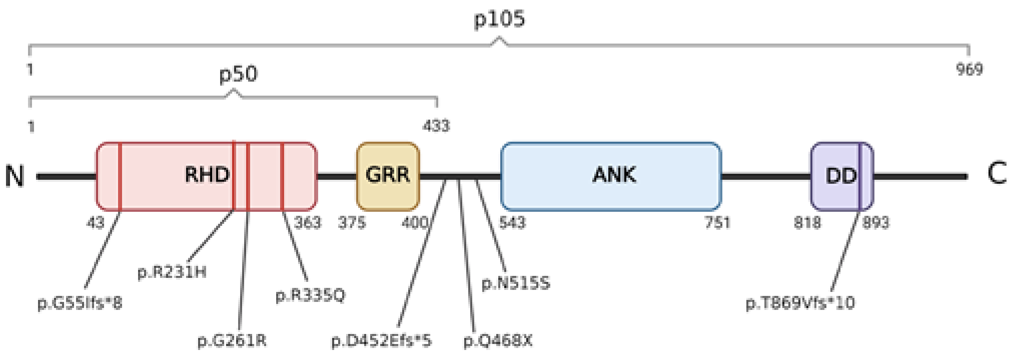


B

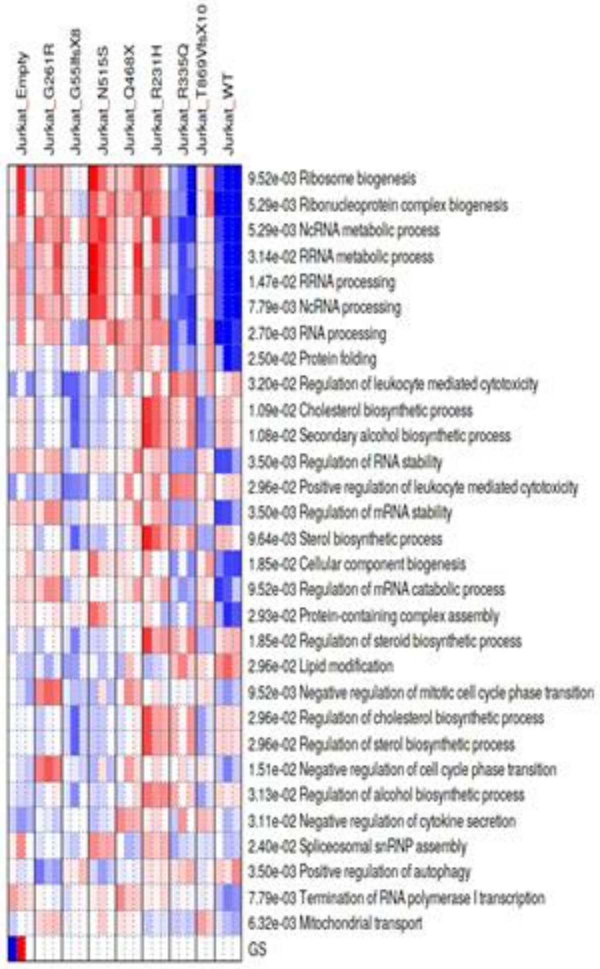


C

Enriched pathways in DEGs for the selected comparison:

| Direction | adj.Pval | nGenes | Pathways |
|---|---|---|---|
| Down regulated | 2.6e-19 | 23 | MSigDB HALLMARK MYC TARGETS V1 |
| | 1.2e-14 | 19 | MSigDB HALLMARK MTORC1 SIGNALING |
| | 3.4e-08 | 13 | MSigDB HALLMARK E2F TARGETS |
| | 7.5e-08 | 10 | MSigDB HALLMARK UNFOLDED PROTEIN RESPONSE |
| | 1.5e-06 | 7 | MSigDB HALLMARK MYC TARGETS V2 |
| | 1.1e-05 | 10 | MSigDB HALLMARK G2M CHECKPOINT |
| | 4.3e-03 | 6 | MSigDB HALLMARK UV RESPONSE UP |
| Up regulated | 4.8e-13 | 21 | MSigDB HALLMARK HYPOXIA |
| | 6.6e-05 | 12 | MSigDB HALLMARK MTORC1 SIGNALING |
| | 1.4e-03 | 10 | MSigDB HALLMARK GLYCOLYSIS |
| | 1.9e-03 | 6 | MSigDB HALLMARK CHOLESTEROL HOMEOSTASIS |
| | 7.0e-03 | 6 | MSigDB HALLMARK ANDROGEN RESPONSE |

D

Enriched pathways in DEGs for the selected comparison:

| Direction | adj.Pval | nGenes | Pathways |
|---|---|---|---|
| Down regulated | 3.9e-14 | 29 | Ribonucleoprotein complex biogenesis |
| | 2.3e-12 | 22 | Ribosome biogenesis |
| | 3.7e-11 | 35 | RNA processing |
| | 5.8e-09 | 25 | NcRNA metabolic process |
| | 6.2e-09 | 16 | RRNA processing |
| | 6.6e-09 | 17 | RRNA metabolic process |
| | 2.0e-08 | 20 | NcRNA processing |
| | 1.6e-05 | 27 | DNA metabolic process |
| | 1.6e-05 | 7 | ER-nucleus signaling pathway |
| | 2.5e-05 | 25 | MRNA metabolic process |
| | 5.1e-05 | 21 | Peptide biosynthetic process |
| | 6.1e-05 | 12 | Protein folding |
| | 6.4e-05 | 23 | Peptide metabolic process |
| | 8.1e-05 | 14 | Regulation of mRNA metabolic process |
| | 8.1e-05 | 39 | Cellular response to stress |

### Supplemental Figure 2

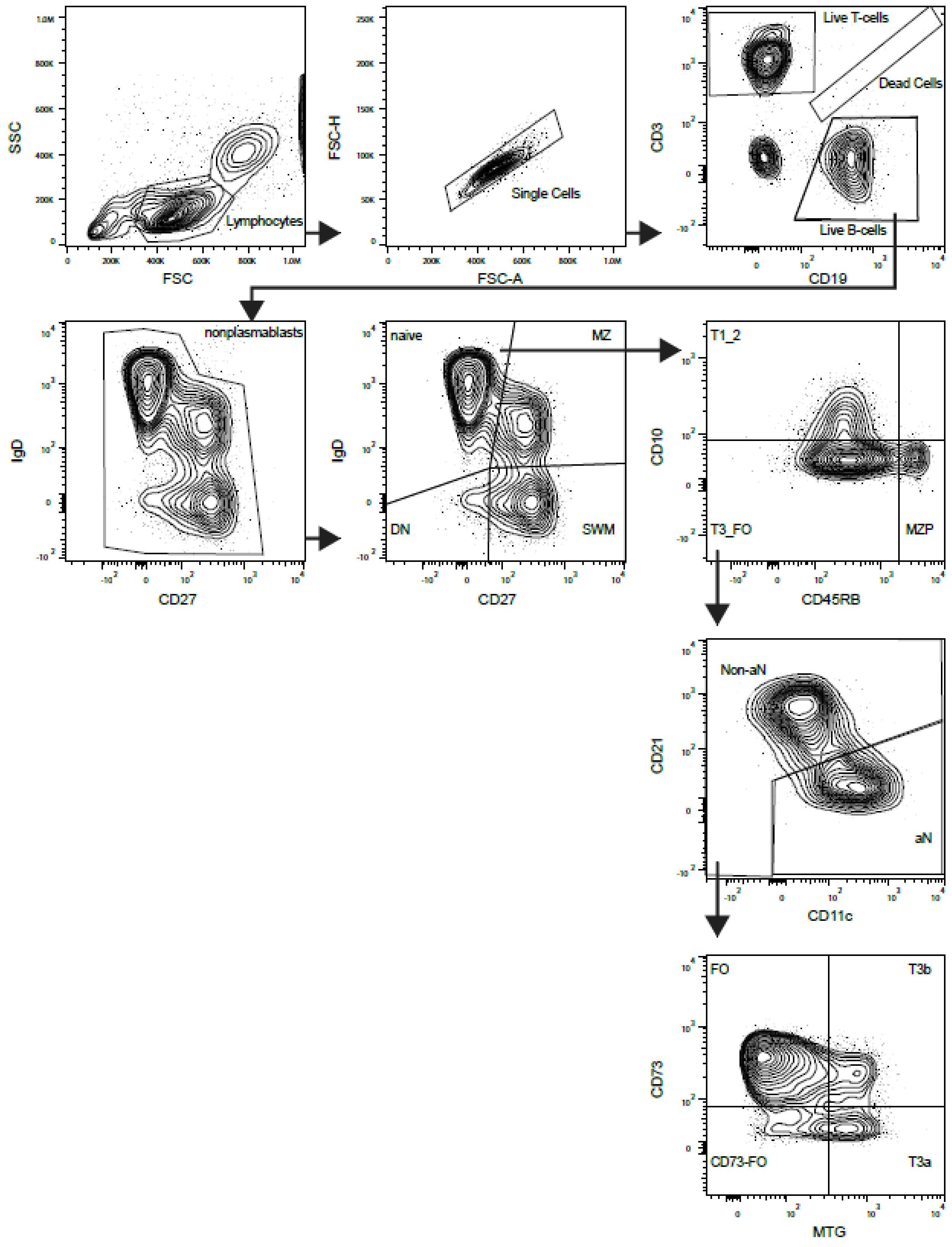

### Supplemental Figure 3

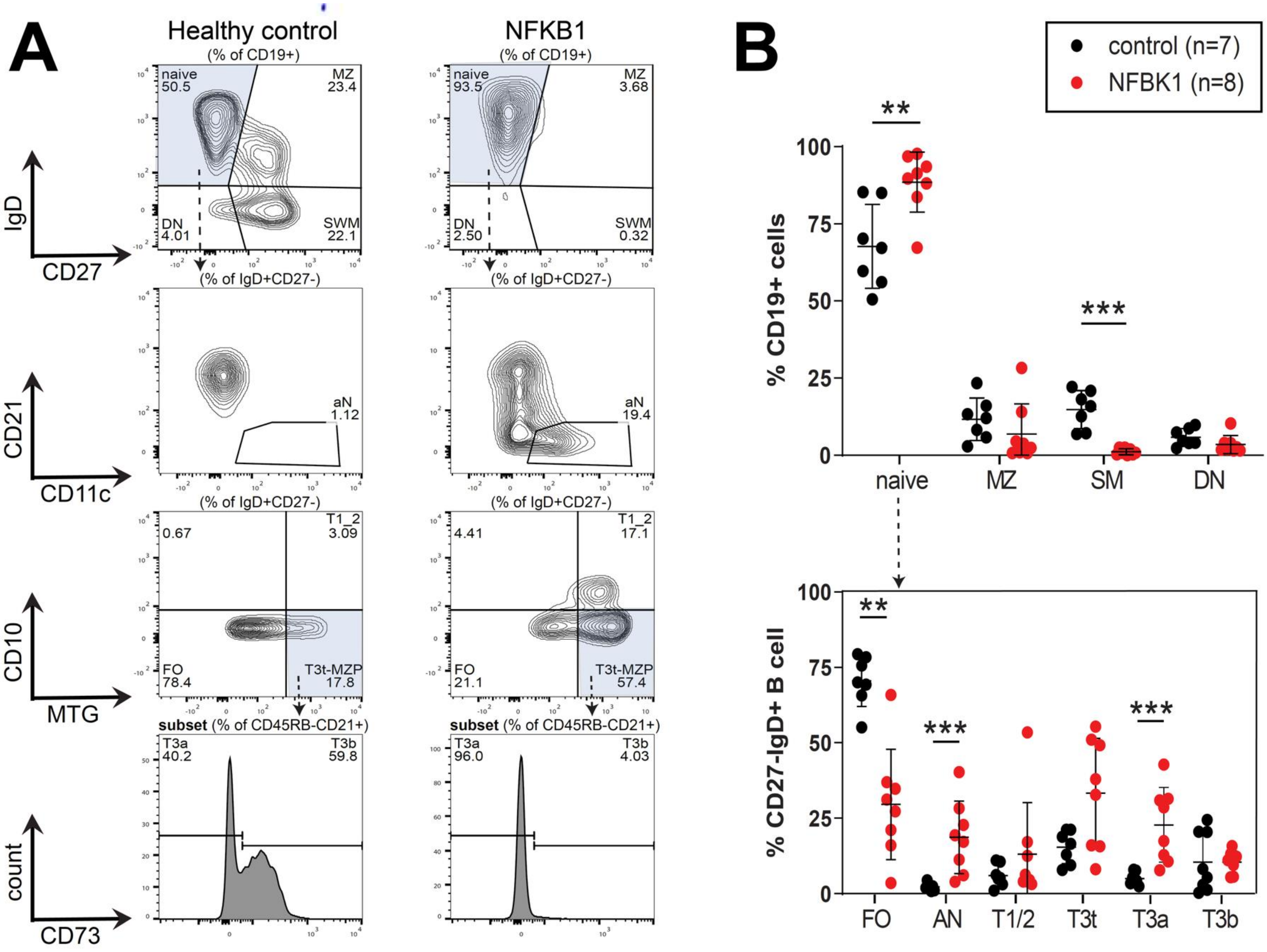

### Supplemental Figure 4

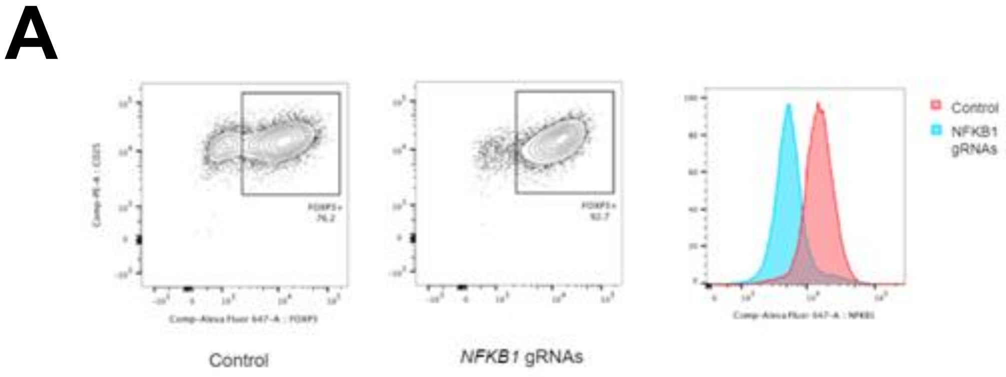


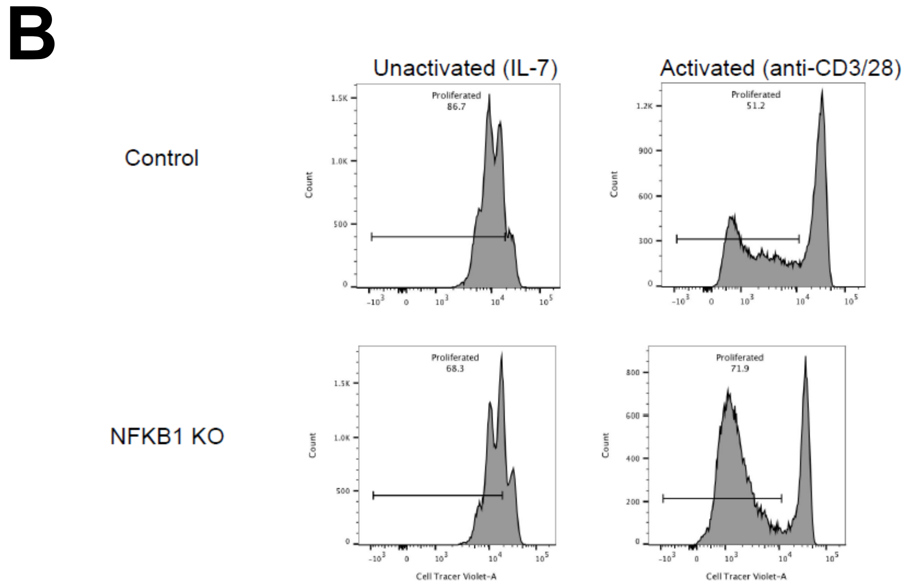

### Supplemental Figure 5

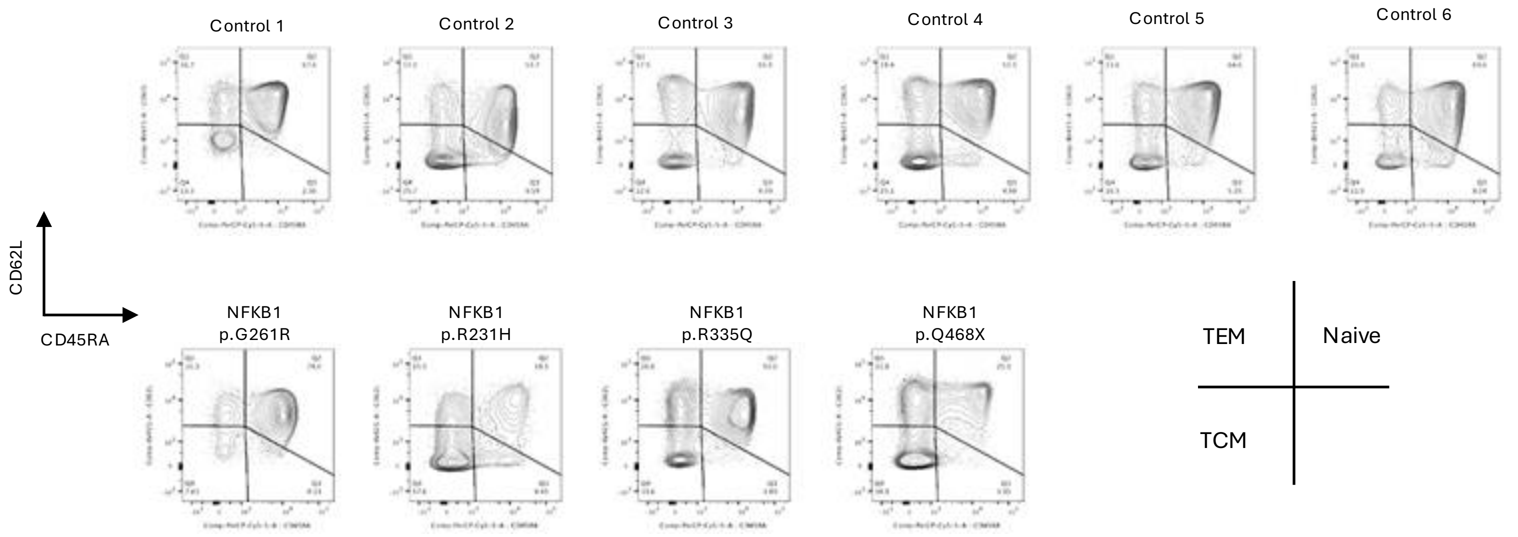

### Supplemental Figure 6

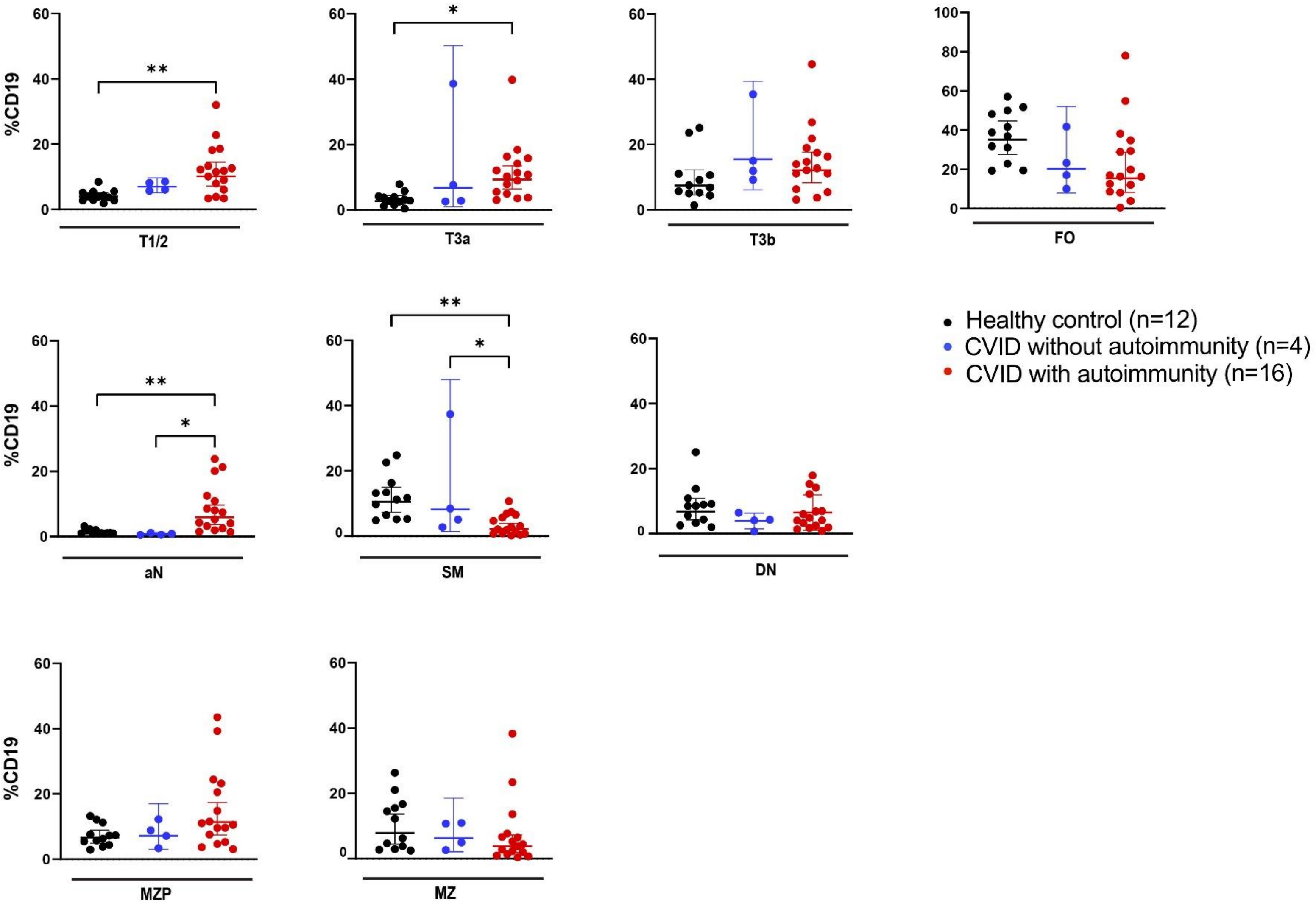

### Supplemental Figure 7

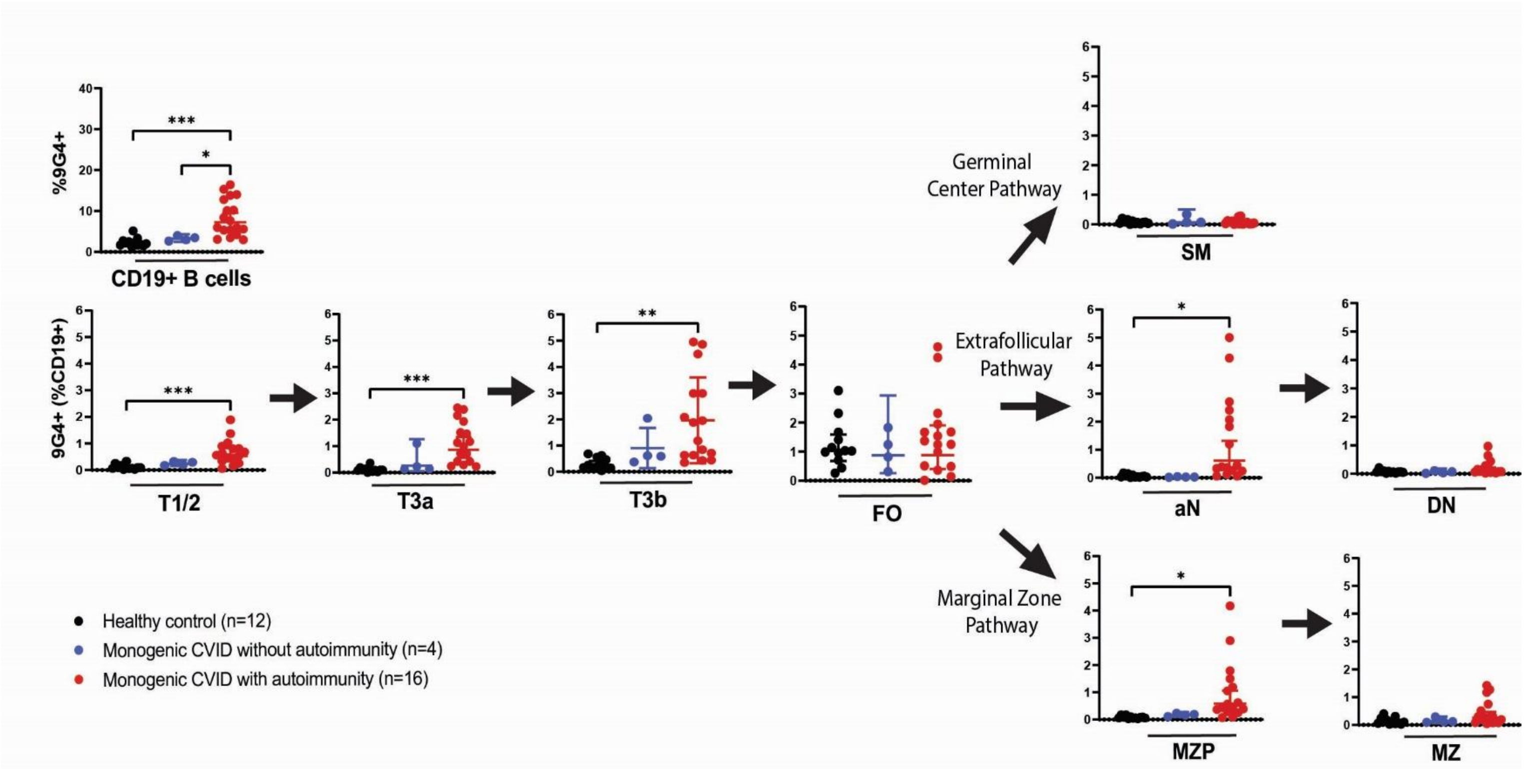

### Supplemental Figure 8

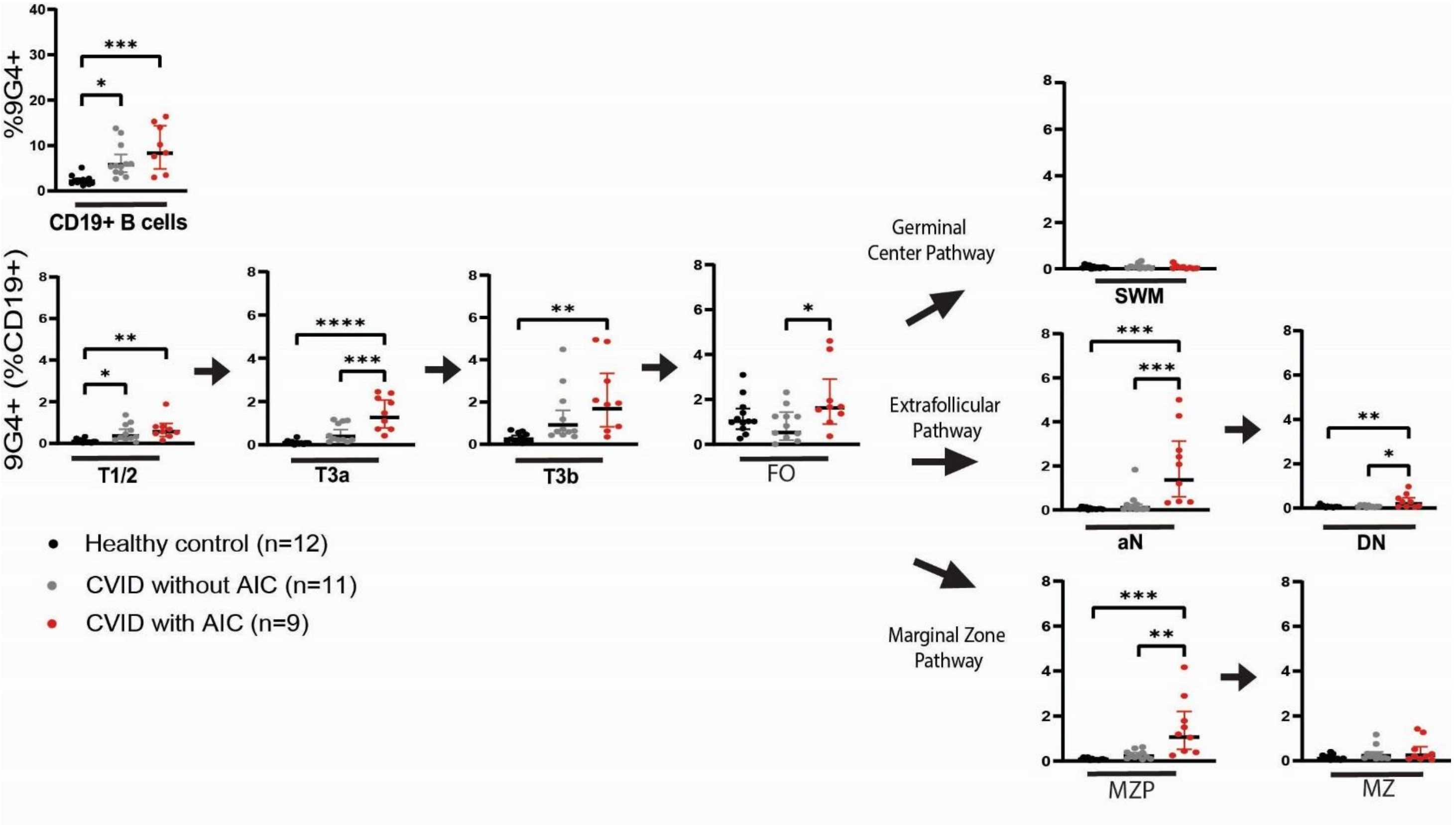

### Supplemental Figure 9

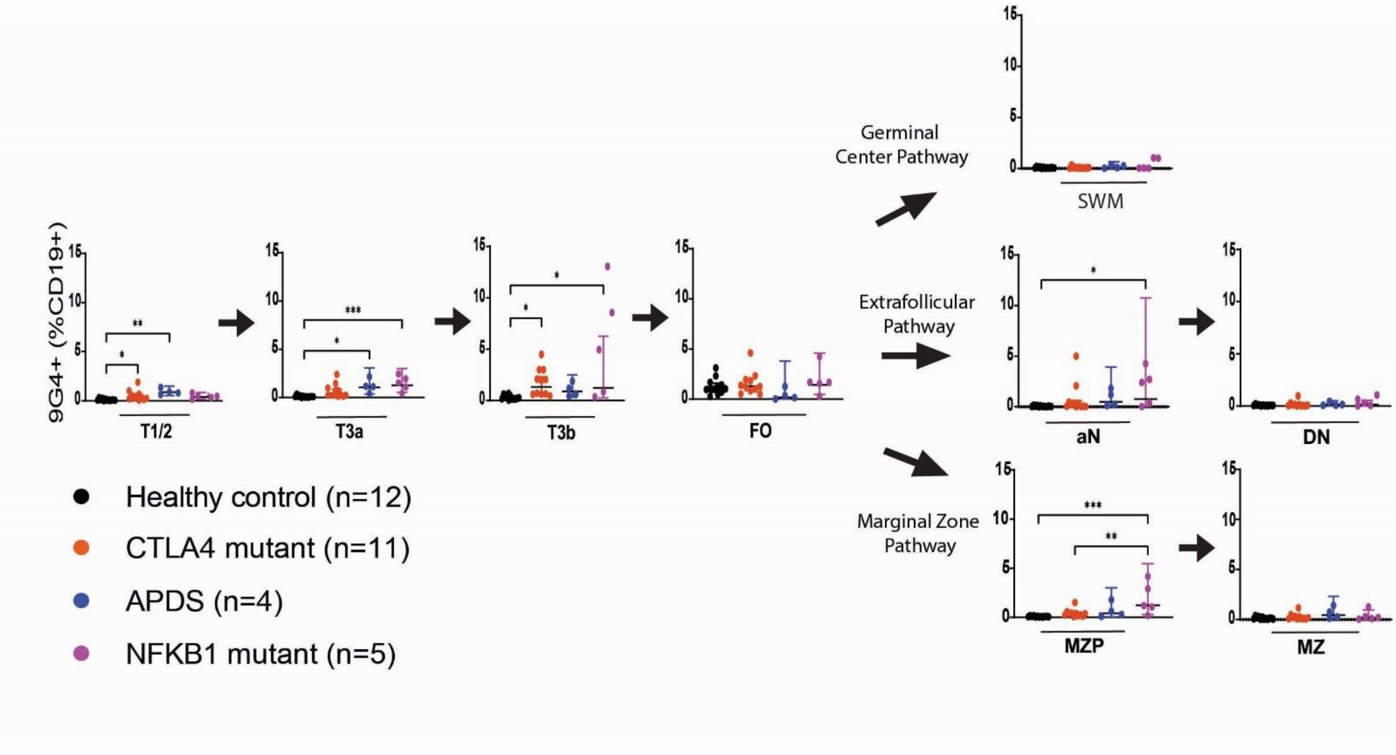

### Supplemental Figure 10

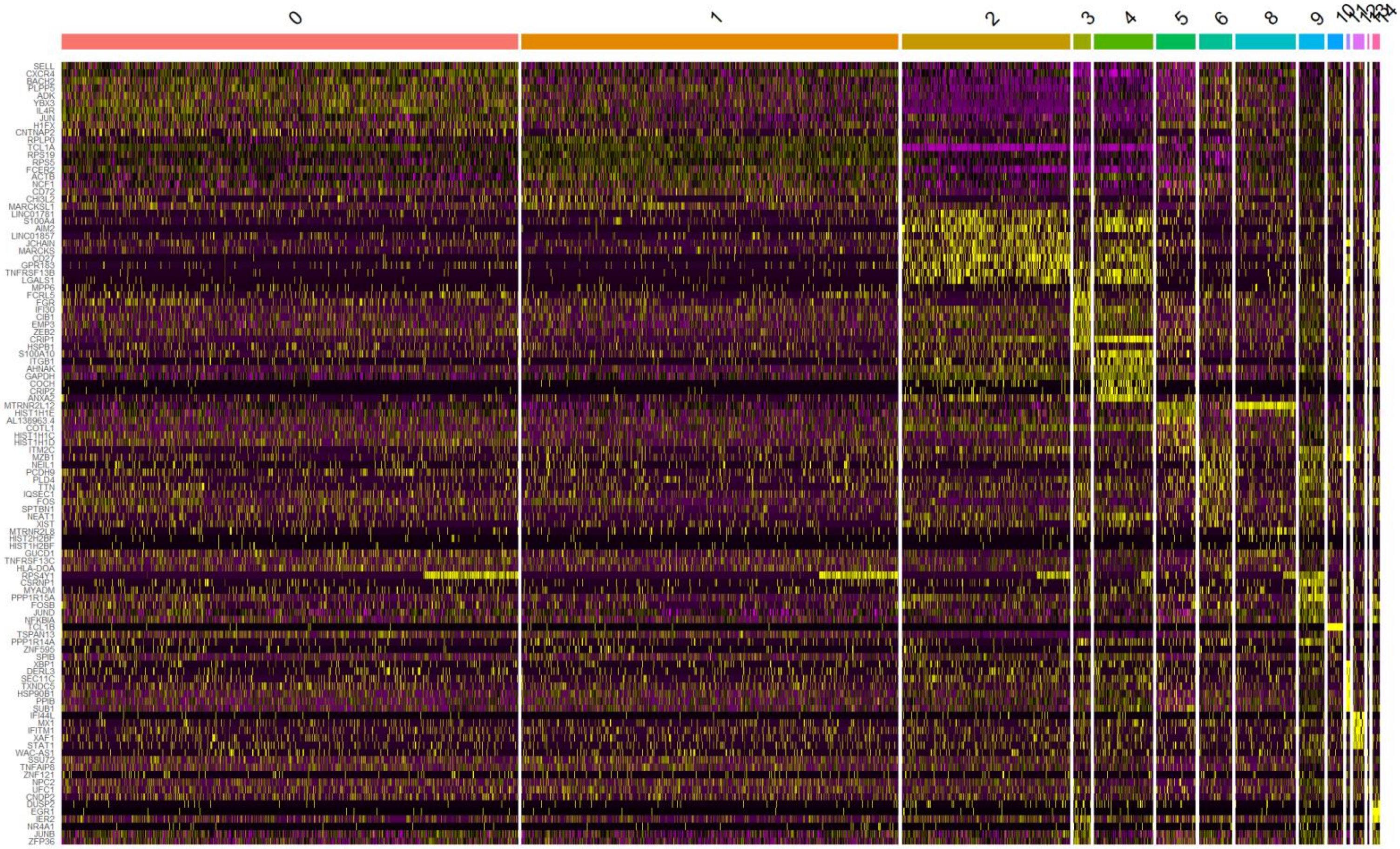

### Supplemental Figure 11

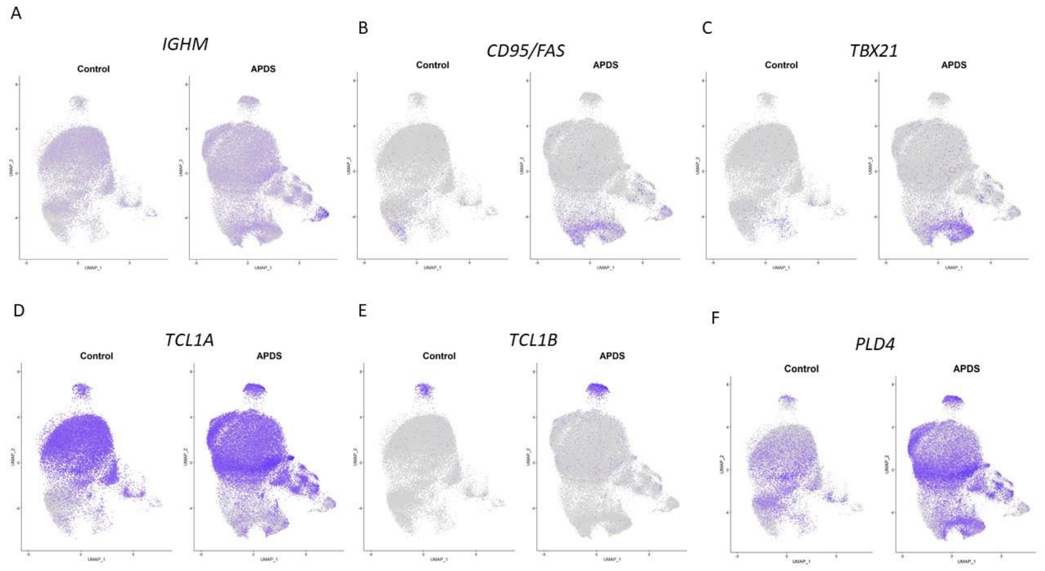

### Supplemental Figure 12

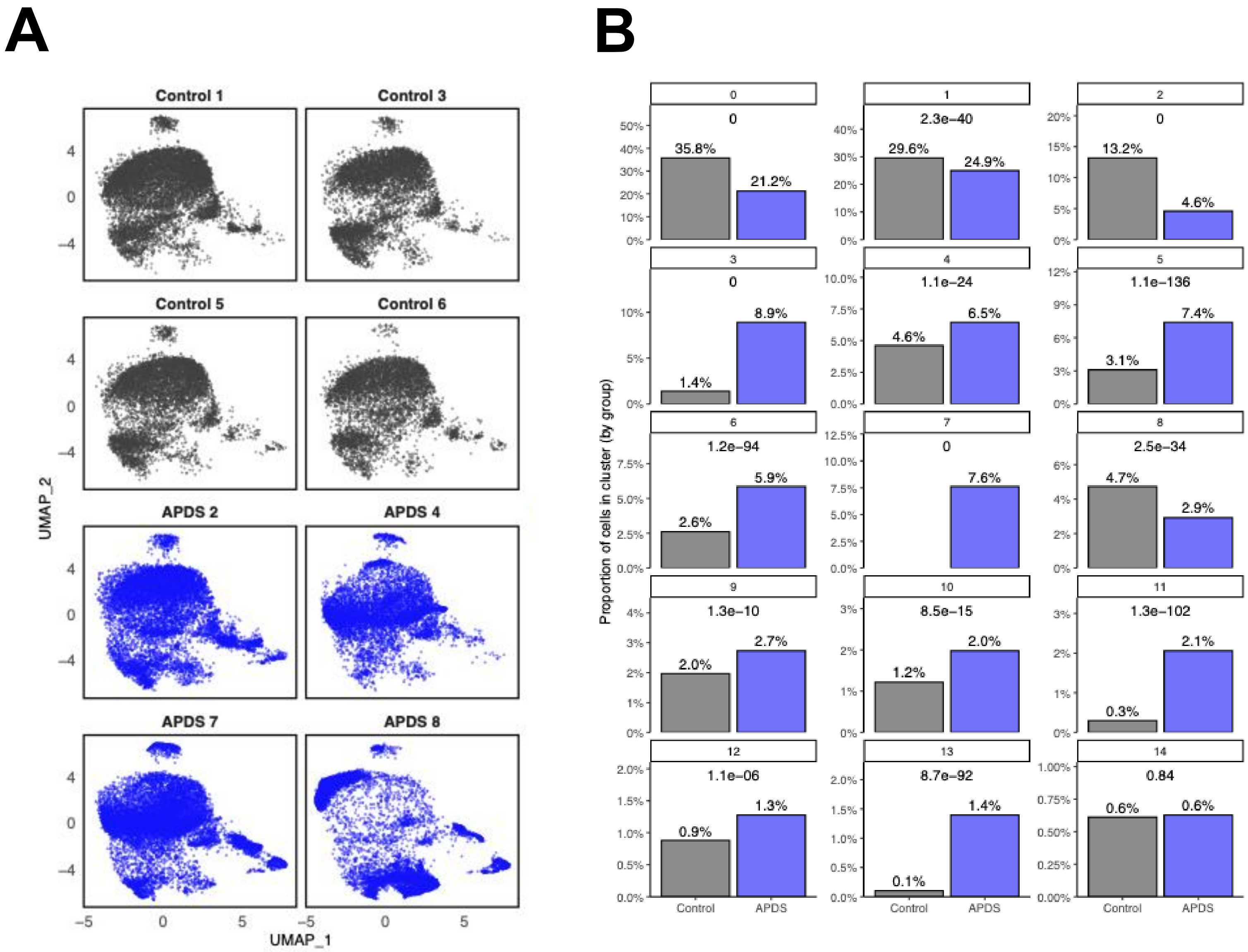

### Supplemental Figure 13

## A

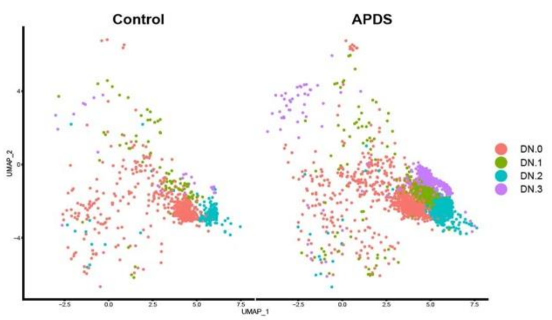


## B

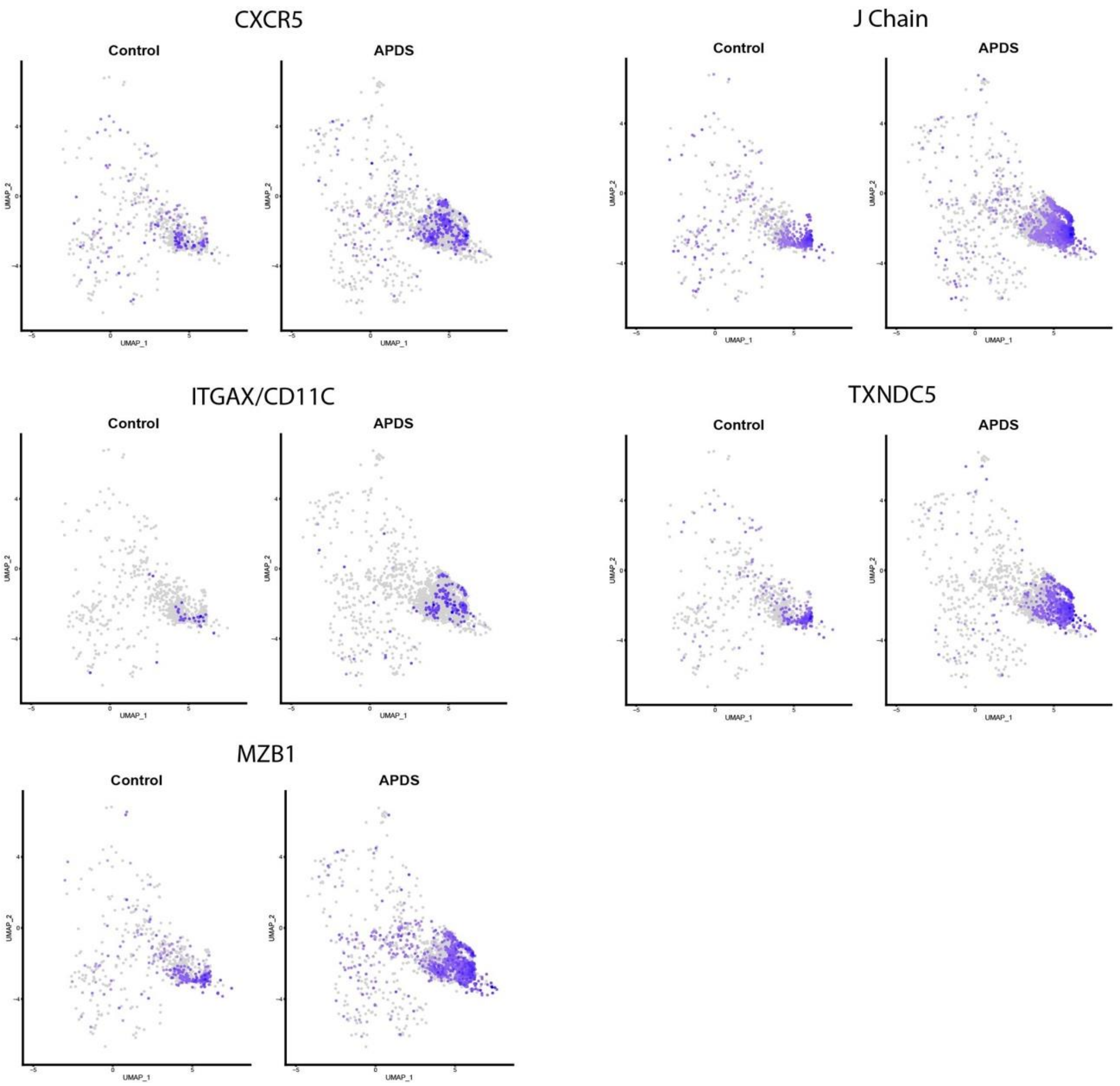

### Supplemental Figure 14

A

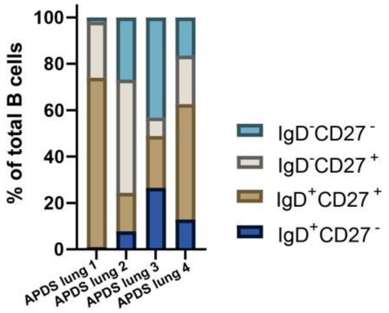


B

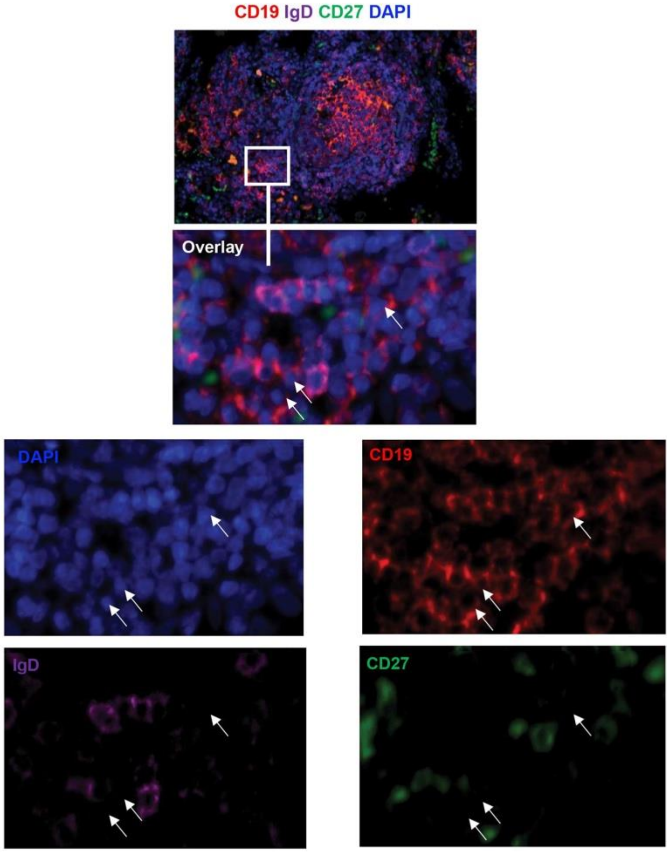

### Table S1. Characteristics of patients with monogenic variants.

| Family | n/a | n/a | n/a | n/a | n/a | n/a | n/a | n/a | n/a | n/a | 4 | 1 | 1 | 1 | 3 | 3 | 3 | 4 | 5 | 5 | 5 |
|---|---|---|---|---|---|---|---|---|---|---|---|---|---|---|---|---|---|---|---|---|---|
| **Member** | n/a | n/a | n/a | n/a | n/a | n/a | n/a | n/a | n/a | n/a | proband | proband | sister | brother | proband | son 1 | son 2 | brother | cousin | son of cousin | proband |
| **Variant; Reference for Functional Studies** | PIK3CD; p.E1021K; [12] | PIK3CD; p.E1021K; [12] | PIK3CD; p.M61V; [12] | PI3KCD; Intron 4, c.371-4C>G | NFKB1 c.1190del (p.Gly397Alafs*35); ClinVar Variation ID: 1352196 | NFKB1 c.1356del, p.Asp452Glufs*5 (het.); Supp Figure 1 | NFKB1; c.692G>A, R231H (het.); Supp Figure 1 | NFKB1; c.2602_2603dupGG, p.T869VfsX10 (het.); Supp Figure 1 | NFKB1; Q468X, (het.); Supp Figure 1 | NFKB1; p.G261R, c.781G>C (het.); Supp Figure 1 | CTLA4; c.81dupT, p.L28SfsX402 (het.); [19] | CTLA4; 3' UTR mutation (homo.); [19] | CTLA4; 3' UTR mutation (homo.); [19] | CTLA4; 3' UTR mutation (het.); [19] | CTLA4; c.410C>T, p.P137L (het.); [19] | CTLA4; c.410C>T, p.P137L (het.); [19] | CTLA4; c.410C>T, p.P137L (het.); [19] | CTLA4; c.81dupT, p.L28SfsX402 (het.); [19] | c.173G>C, CTLA4; p.C58S (het.); [19] | CTLA4; c.173G>C, p.C58S (het.); [19] | CTLA4; c.173G>C, p.C58S (het.); [19] |
| **Other Genetic Variants** | None | None | None by GeneDx | Invitae, Gene Results of Uncertain Significance: ARPC1B, COL7A1, DEF6, IL12RB1, PIK3CD, TONSL | Invitae, Gene Panel (574 genes tested), no other leading variants | Invitae, EPG5 c.290C>T (p.Thr97Met) heterozygous, EXTL3 c.724G>T (p.Val242Leu), KMT2D c.1954C>T (p.Arg652Cys), MKL1 c.1694_1695insACCCGC (p.Ala569_Pro570dup), PTPRC c.1297G>A (p.Asp433Asn), UNC13D c.334T>C (p.Cys112Arg) heterozygous | None | None | None | None | *TNFRSF13B* (p.A181E) *NOD2* (p.L1007PfaX2), ADA2 (deletion exon 7) | None | None | None | None | None | None | ADA2 (deletion exon 7) | None, CTLA4 tested via family-variant testing program | CHD7 | Negative Invitae ALPS gene panel |
| **Age at B cell immunophenotyping** | 28 years | 47 years | 28 years | 34 years | 34 years | 27 years | 56 years | 69 years | 64 years | 23 months | 13 years | 23 years | 25 years | 31 years | 45 years | 20 years | 17 years | 8 years | 49 years | 13 years | 51 years |
| **Gender** | Male | Female | Male | Female | Female | Male | Male | Female | Male | Male | Male | Male | Female | Male | Female | Male | Male | Male | Female | Male | Female |
| **Lymphoproliferation** | LAD, AIE Splenomegaly | LAD, GLILD Splenomegaly | LAD, GLILD, AIE, Splenomegaly | None | GLILD, LAD, Splenomegaly | Splenomegaly | NRH, GLILD, splenomegaly, LAD | NRH, AIE, GLILD, NLH of gastric body mucosa, splenomegaly, LAD, skin cancer | Prostate cancer | None | LAD, Splenomegaly | GLILD, NRH, splenomegaly, LAD | None | n/a | Splenomegaly | n/a | LAD | n/a | GLILD, sarcoid, LAD | Asthma | GLILD, LAD |
| **Autoimmunity** | AIC – no; AI other - Celiac | AIC – no; AI other - no | AIC - ITP, AIHA, chronic neutropenia; AI other - no | AIC – no; AI other - ANCA+ granulomatosis with polyangitis | AIC – Evans Syndrome; AI other - Psoriasis | AIC – ITP, neutropenia; AI other - Alopecia, vitiligo, + lupus anticoagulant | AIC – AIHA, ITP; AI other - AIH, IBD, DM | AIC – AIHA, ITP; AI other – AIH | AIC – ITP; AI other - no | AIC – no; AI other - Myositis | AIC – no; AI other - Colitis | AIC – AIHA, ITP; AI other - DM, AIE | AIC – ITP; AI other: severe RA (Seronegative inflammatory arthritis) | n/a | AIC - no; AI other - Hashimoto's thyroiditis, seronegative inflammatory arthritis | n/a | n/a | n/a | AIC – ITP; AI other - inflammatory arthritis, AIE, hypothyroidism | AIC - no; AI other – psoriasis, chronic diarrhea | AIC - ITP, AIHA; AI other- AIE |
| **Bronchiectasis** | + | + | + | - | - | - | + | + | + | - | - | + | - | n/a | - | n/a | - | - | - | - | - |
| **ALC (1000-4800 cells/µL)** | 570 | 1770 | 270 | n/a | 910 | n/a | 376 | 1487 | 889 | 3282 | 1166 | n/a | n/a | n/a | n/a | n/a | n/a | 2122 | 955 | 3172 | 1060 |
| **CD3+ (690-2540 cells/µL)** | 731 | 1434 | 495 | 2311 | 863 | 683 | 376 | 1487 | 889 | 3282 | 791 | 275 | 1077 | n/a | 255 | n/a | 955 | 1457 | 762 | 2208 | n/a |
| **CD4+ (419-1590 cells/µL)** | 260 | n/a | 211 | 768 | 480 | 316 | 169 | 1078 | 474 | 2067 | 357 | 160 | 642 | n/a | 139 | n/a | 434 | 940 | 598 | 1324 | n/a |
| **CD4+CD45RA+ (% CD4+)** | 6.1 | n/a | 3.0 | 23.6 | 8.6 | 15.2 | 12.9 | 0.9 | 17.8 | 74.4 | n/a | 1.7 | n/a | n/a | 1.1 | n/a | 24.4 | n/a | 6 | 34 | n/a |
| **CD4+CD45RO+ (% CD4+)** | 89.5 | n/a | 87.8 | 59 | n/a | 83.2 | 83 | 94.5 | 67.67 | 16.1 | n/a | 97.1 | n/a | n/a | 87.4 | n/a | 67.5 | n/a | n/a | n/a | n/a |
| **CD8+ (190-1140 cells/µL))** | 439 | n/a | 259 | 1452 | 394 | 316 | 215 | 419 | 392 | 965 | 378 | 112 | 400 | n/a | 107 | n/a | 366 | 454 | 119 | 745 | n/a |
| **CD8+CD45RA+ (% CD8+)** | 58.6 | n/a | 57.6 | 67 | 16.8 | 70.7 | 47.1 | 31 | 63 | 89.5 | n/a | 22.4 | n/a | n/a | 11.6 | n/a | 42.5 | n/a | 5 | 24 | n/a |
| **CD8+CD45RO+ (% CD8+)** | 29.3 | n/a | 15.4 | 27.7 | n/a | 28.2 | 44.7 | 41.5 | 25.3 | 5.1 | n/a | 68.8 | n/a | n/a | 60.6 | n/a | 54.1 | n/a | n/a | n/a | n/a |
| **CD19+ (90-660 cells/µL)** | 121 | 142 | 77 | 749 | 19 | 118 | 87 | 324 | 129 | 1083 | 272 | 14 | 86 | n/a | 33 | n/a | 484 | 404 | 114 | 727 | n/a |
| **CD3-CD16/56+ (90-590 cells/µL)** | 174 | 160 | 40 | 140 | 26 | 80 | 73 | 274 | 433 | 245 | 64 | 55 | 122 | n/a | 128 | n/a | 208 | 206 | 78 | 214 | n/a |
| **CD27+ (% CD19+)** | 8.9 | 20.5 | 13.3 | 7 | 33.3 | 9.3 | 2.3 | 2.10 | 23.6 | 10.1 | 24 | 4 | 5 | n/a | 22.1 | n/a | 10.3 | 19 | 12 | 23 | n/a |
| **CD27+IgD/M-** | 0.8 | 6.0 | 0.1 | 2 | 0.9 | 1.3 | <0.1 | 0.30 | <0.1 | 3.9 | n/a | 0.4 | <1 | n/a | 4 | n/a | 1.7 | n/a | n/a | n/a | n/a |

| | | | | | | | | | | | | | | | | | | | | | |
|---|---|---|---|---|---|---|---|---|---|---|---|---|---|---|---|---|---|---|---|---|---|
| **(% CD19+)** | | | | | | | | | | | | | | | | | | | | | |
| **IgG (614-1295 mg/dL)** | 969 | 445 | 87 | 243 | 591 (on IVIG) | 537 | 33 | 1337 | <40 | Not measured before IVIG | 748 | 422 | 696 | n/a | 567 | n/a | 648 | 662 | 512 | 979 | 544 |
| **IgA (69-309 mg/dL)** | 64 | 5 | 13 | <7 | <26 | 96 | 7 | <7 | <7 | 87 | 59 | 7 | 95 | n/a | 38 | n/a | 52 | 105 | 11 | 84 | 167 |
| **IgM (53-334 mg/mL)** | 284 | 641 | 388 | 930 | <19 | 30 | 13 | 29 | <5 | 295 | 15 | 5 | 358 | n/a | 66 | n/a | 71 | 14 | 52 | 62 | 143 |
| **Pneumococcal titers (≥ 1.3 μg/mL)** | 10/23+ (post-pneumovax) | 0/14+ (post-pneumovax) | 0/23+ (post-pneumovax) | 5/23+ (post-pneumovax) | n/a | 18/23+ (post-pnuemocax) | n/a | n/a | n/a | 18/23 (post-pneumovax) | 0/23+ (post-pneumovax) | n/a | 2 of 23 serotypes (8.7%) | n/a | 82.6% | n/a | n/a | 0 of 14→8 of 14 In 2013 | 2 of 23 | 4 of 23→21/23 | n/a |
| **Proliferation to PHA** | n/a | n/a | Decreased | Normal | n/a | Normal | Decreased | Normal | Normal | Normal | Normal | Decreased | Normal | n/a | n/a | n/a | n/a | n/a | Normal (high) | Normal | n/a |
| **Proliferation to PWM** | n/a | n/a | Insufficient | Normal | n/a | Normal | Normal | Normal | Normal | Normal | Normal | Insufficient | Insufficient | n/a | n/a | n/a | n/a | n/a | Normal | Normal | n/a |
| **Proliferation to Candida** | n/a | n/a | Insufficient | Normal | n/a | Normal | Normal | n/a | Normal | Decreased | Normal | Insufficient | Normal | n/a | n/a | n/a | n/a | n/a | Normal | Normal | n/a |
| **Proliferation to Tetanus** | n/a | n/a | Insufficient | Decreased | n/a | Decreased | Decreased | n/a | Normal | Decreased | Insufficient | Insufficient | Normal | n/a | Normal | n/a | n/a | n/a | Normal | Normal | n/a |
| **Proliferation to aCD3/IL2** | n/a | n/a | Not sent | n/a | n/a | Normal | n/a | Normal | n/a | Normal | n/a | Normal | n/a | n/a | n/a | n/a | n/a | n/a | n/a | n/a | n/a |
| **Treatment prior to B cell phenotyping** | SCIG | IVIG | SCIG; rituximab | None | None | | None | None | None | None | IVIG | IVIG | IVIG (Tofacitinib starting in 2016) | None | Prednisone | None | None | n/a | n/a | n/a | Dexamethasone, Prednisone, IVIG, Rituximab, Romiplostim |

Normal reference ranges from the Massachusetts General Hospital shown where applicable. Pneumococcal titers shown as ratio of positive over tested serotypes. Autoimmune enteropathy (AIE); autoimmune hepatitis (AIH); autoimmune hemolytic anemia (AIHA); diabetes mellitus (DM); immune thrombocytopenia (ITP); intravenous immunoglobulin (IVIG); granulomatous-lymphointerstitial lung disease (GLILD); human papilloma virus (HPV); lymphadenopathy (LAD); nodular regenerative hyperplasia (NRH), not available (n/a); rheumatoid arthritis (RA); subcutaneous immunoglobulin (SCIG); varicella zoster virus (VZV).

| Cluster Number | Cluster Short Name | Designation | CITE-seq surface marker | RNA differential gene expression | Gene set enrichment analysis OXPHOS pathway | APDS patient cluster expansion |
|---|---|---|---|---|---|---|
| 0 | FO | Follicular | $IgD^{+}10^{-}$ | N/A | Low | N/A |
| 1 | T3b | Transitional T3b | $IgD^{+}CD10^{+/-}$ | IL-4R HI | Low (Controls), High (APDS) | Expanded in APDS |
| 2 | MZ | Marginal Zone | $IgD^{lo}CD10^{-}$ | $CD27^{+}IgM^{hi}PL4^{+}CD1c^{+}YBX1^{hi}$ | High | N/A |
| 3 | aN | Activated naïve | $IgD^{+}$ | $CD11c^{+}FAS^{+}TBX21^{+}$ | High | Expanded in APDS |
| 4 | SWM | Switched memory | $IgD^{-}$ | $CD27^{+}CXCR5^{+}$ | High | N/A |
| 5 | DN1/DN4 | Double negative DN1/DN4 | $IgD^{lo/-}CD10^{-}$, few $CD11c^{+}$ | N/A | Low | Expanded in APDS |
| 6 | T1/2a | Transitional T1/2 activated | $IgD^{+}CD10^{hi}$ | $CD38^{hi}$ | Low | Expanded in APDS |
| 7 | aT1/2 | Activated Transitional T1/2 | $IgD^{+}CD10^{hi}$ | $CD38^{hi}$ | High | Unique to APDS |
| 8 | MZP | Marginal Zone Progenitors | $IgD^{+}CD10^{-}$ | $CD27^{-}CD38^{mid}CD24^{mid}$ | Low (Controls), High (APDS) | N/A |
| 9 | T1/2 | Transitional T1/2 | $IgD^{+}CD10^{hi}$ | $CD38^{hi}$ | High | N/A |
| 10 | $TCL1B^{+}T3b$ | $TCL1B^{+}$ Transitional T3B | $IgD^{+}CD10^{lo}$ | $CD10^{hi}TCL1B^{++}$ | Low (Controls), High (APDS) | Expanded in APDS |
| 11 | PB | Plasmablasts | N/A | $JChain^{hi}MZB1^{hi}XBP\text{-}1^{+}UPR^{hi}$ | High | Expanded in APDS |
| 12 | $IFN^{+}T3a$ | IFN activated Transitional T3a | $IgD^{+}CD10^{lo}$ | N/A | Low (Controls), High (APDS), +IFN signaling | N/A |
| 13 | DN2/DN3 | Double negative DN2/DN3 | $IgD^{-}CD10^{lo}$ | $CD27^{lo}$ | Low | Expanded in APDS |
| 14 | ABC-like | Age-associated B cells | $IgD^{-}$ | $CD27^{+}EGR1^{+}AHNAK^{+}CD11c^{+}$ | N/A | N/A |

**Table S2: List of B cell cluster designations.** Annotated by single cell CITE-seq surface marker expression, single-cell RNA-sequencing differential gene expression, and gene set enrichment analysis for oxidative phosphorylation pathway. Final column designates if cluster is expanded in APDS patients compared to healthy controls.
